# Single molecule localization microscopy with autonomous feedback loops for ultrahigh precision

**DOI:** 10.1101/487728

**Authors:** Simao Coelho, Jongho Baek, Matthew S. Graus, James M. Halstead, Philip R. Nicovich, Kristen Feher, Hetvi Gandhi, Katharina Gaus

## Abstract

Single-molecule localization microscopy (SMLM) promises to provide truly molecular scale images of biological specimens^1–5^. However, mechanical instabilities in the instrument, readout errors and sample drift constitute significant challenges and severely limit both the useable data acquisition length and the localization accuracy of single molecule emitters^6^. Here, we developed an actively stabilized total internal fluorescence (TIRF) microscope that performs 3D real-time drift corrections and achieves a stability of ≤1 nm. Self-alignment of the emission light path and corrections of readout errors of the camera automate channel alignment and ensure localization precisions of 1-4 nm in DNA origami structures and cells for different labels. We used Feedback SMLM to measure the separation distance of signaling receptors and phosphatases in T cells. Thus, an improved SMLM enables direct distance measurements between molecules in intact cells on the scale between 1-20 nm, potentially replacing Förster resonance energy transfer (FRET) to quantify molecular interactions^7^. In summary, by overcoming the major bottlenecks in SMLM imaging, it is possible to generate molecular images with nanometer accuracy and conduct distance measurements on the biological relevant length scales.

---

Improvement in optics and fluorophore design have led to super-resolution methods such as (direct) Stochastic Optical Reconstruction Microscopy ((d)STORM)^2^ and DNA point accumulation for imaging in nanoscale topography (DNA-PAINT)^3^ that are anticipated to revolutionize biology as they can localize individual molecules in the cellular context. In SMLM, single fluorophores are temporally separated so that a sparse set of point emitters can be captured in each camera frame. The temporal separation is achieved via photoactivation^1^, stochastic switching of fluorophores between a fluorescent and non-fluorescent state^2^ or the reversible binding of fluorophore to the target site^3^. The sequential imaging of individual fluorophores, however, requires long acquisition times as typically tens of thousands of frames are needed to map a given protein in a cell. In theory, the localization precision of each fluorophore only depends on the number of photons collected at each point. However, drift during sample acquisition makes it challenging to correctly assign photons to the fluorescent molecule and reduces the localization precisions to tens of nanometers^6, 8^. Previous attempts to achieve ultrahigh resolution therefore had to rely on averaging a large number of frames^9^ or could only detect the subset of fluorophores that emitted an unusual high number of photons^10, 11^.

Most current strategies to compensate for drift in SMLM use post-acquisition corrections where the position of individual molecules (and their localization precision) is first calculated prior to the drift correction. The average displacement of multiple fiducial markers placed in the sample are used to correct the spatial coordinates of the localizations detected, on a frame-by-frame basis^12^. Post-acquisition corrections do not improve the localization precision of a single emission and make long data acquisitions challenging, for example for sequential multi-color DNA-PAINT imaging. This is because most post-acquisition drift corrections cannot correct for drift greater than 1 pixel (~100 nm). Thus, drift and vibration during data acquisition are major limiting factors for SMLM, reducing both localization accuracy and acquisition length (Fig. 1a and Supplementary Fig. 1).

**Figure 1:**
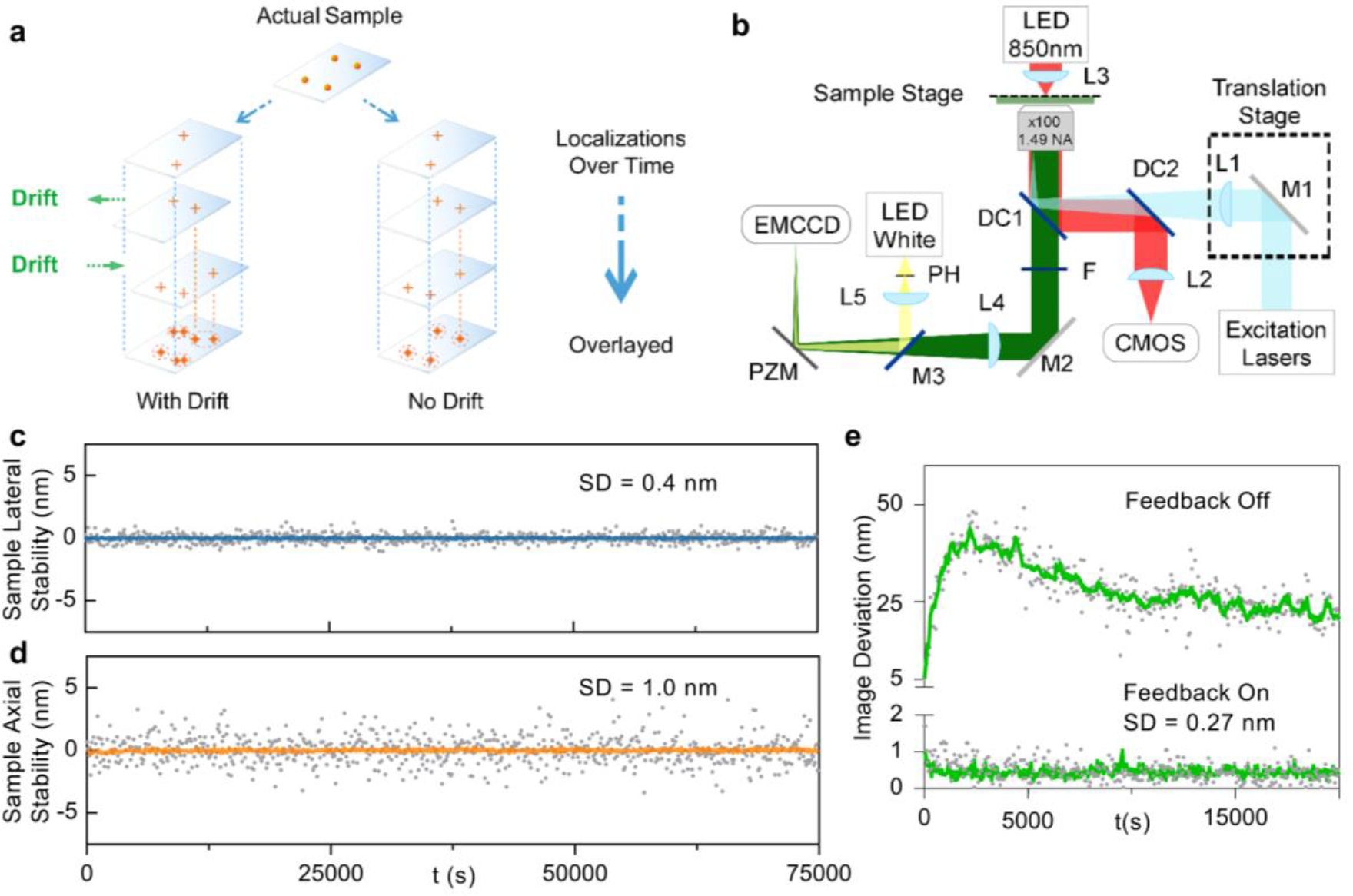
Feedback SMLM. **a**, Principle of active stabilization for SMLM. Drift degrades the localization precision of single emitters. Active stabilization is performed at faster rates than the on-time of the molecules, thus aligning photons from a single emitter within a single frame. **b**, Optical Setup. Fluorescence, excited by lasers (light blue) under TIRF illumination (via a translation stage), is collected and imaged onto an EMCCD camera. In addition, a white LED is combined with the emission light path and produces a reference point for the piezoelectric feedback correction. Sample drift is determined with 3 μm polystyrene fiducial beads in the sample, which are illuminated with an infrared LED and imaged with a dedicated CMOS camera. Sample drift is corrected in real-time at a frequency of 10-15 Hz. M1-2, mirrors; M3, 0.5% reflective mirror; L1-5, lenses; F, filter; DC1-2, dichroics; PZM, piezoelectric mirror; PH, pinhole; EMCCD, EMCCD camera. In **c** and **d**, the lateral and axial stability of the sample is reflected in the standard deviations of 0.4 nm and 1 nm, respectively. **e**, Image deviation without the feedback is 45 nm and with the feedback is 0.27 nm. In c-e, grey symbols represent sample and image deviation; blue, orange, green lines represent a 10-point average (1/1000 points plotted).

We have developed an SMLM system that performs real-time drift correction in 3D (Fig. 1b). The approach realigns molecular emissions independently from the fluorescence, therefore readjusting the point-spread function (PSF) of the molecules while these are still emitting. The system, termed Feedback SMLM, uses three types of corrections. First, there is a feedback loop between the sample and stage position (Supplementary Fig. 2). A key feature is that non-fluorescent fiducials are used outside the field of view (FoV) of the cameras for SMLM, providing great flexibility in sample positioning and ensuring that fiducials do not interfere with data acquisition (Supplementary Fig. 3). Thus, stage corrections can occur during the acquisition of fluorescence data and are completely independent of the sample nature and density of fluorophores. The non-fluorescent fiducials are 3 μm polystyrene beads that are illuminated with an infrared LED to create diffraction rings. The interference pattern is imaged onto a separate CMOS camera (112 μm × 70 μm FoV, ~5-fold larger than the EMCCD camera) at a maximum speed of 370 frames/s. The diffraction rings of the 3 μm beads provide information of both the *x/y* and *z* position^13–15^. The position is calculated sufficiently fast to facilitate stage corrections at 10-15 Hz (Supplementary Figs. 4 and 5). The active stage correction provides a stabilization of 0.4 nm and 1 nm (standard deviation) in the lateral (*x*/*y*) and axial (*z*) directions, respectively, over hours (Fig. 1c, d). Indeed, unlike in other SMLM instruments, stage drift is independent of acquisition period or imaging parameters (i.e. exposure time). A further advantage of our autonomous optical feedback loop for stage positioning is that the sample position is maintained during buffer exchanges, for example during multi-color DNA-PAINT experiments (Supplementary Fig. 6).

Second, in addition to the stage/sample stabilization, we also use an autonomous optical feedback loop in the emission path. A white LED creates an optical fiducial on the EMCCD camera, which is localized with a precision of 0.05 nm. The piezo-electric mirror is used to account for drift of the optical components and channel alignment. The image drift is reduced from 45 nm to 0.27 nm (Fig. 1e). The third and final correction reduces the variation registered across the EMCCD (Supplementary Fig. 7).

Our technique is particularly useful for prolonged acquisitions such as DNA-PAINT, where improved precision can be achieved via repeated binding events to the same docking site^3^. We demonstrate ultrahigh resolution (~1 nm localization precision) of individual fluorescent accumulation points with DNA origami structures that consist of 3 docking sites which are separated by 20 nm (Fig. 2a, b). Even without averaging, summation, individual correction of binding sites, or any assumption regarding the number of docking sites or their separation, the reconstruction of a single origami ruler is possible. We also demonstrate the improvement in resolution of the Feedback SMLM over standard SMLM imaging without active drift correction (Fig. 2c, d). The standard SMLM images were acquired with gold nanorods on the coverslip and data were drift-corrected post-acquisition using redundant cross-correlation (RCC), a high-precision drift correction method^16^ (Supplementary Figs. 8 and 9).

**Figure 2:**
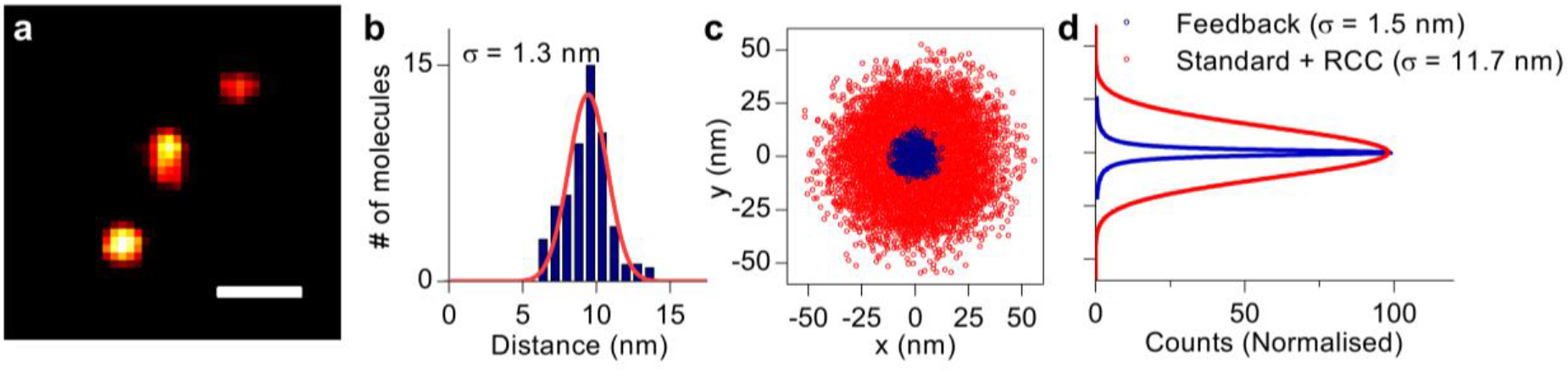
Feedback SMLM imaging of DNA origamis. **a**, DNA-PAINT image of a single origami ruler. Scale bar = 20 nm. **b**, Cross-sectional profile of a single binding site. Red line represents Gaussian fit. **c**, xy distributions of localization points for individual binding sites. The distributions for each binding site was aligned by their respective centre and superimposed. **d**, Cross-sectional fits of (c). **c** and **d**, Blue symbols and lines represent data from Feedback SMLM; red symbols and lines represent data from standard SMLM with post-acquisition redundant cross-correlation (RCC).

We next used the Feedback SMLM to image F-actin (Fig. 3 and Supplementary Fig. 10), an abundant protein that organizes into various structures^17^. Even without post-acquisition processing or customized filament analysis^18^, it was possible to identify single filaments and determine their widths as 5-9 nm without averaging (Fig. 3D, Supplementary Figs. 10 and 11). Grouping of fluorescent events which belong to the same molecule as frequently done in (d)STORM, would further improve the localization accuracy. Furthermore, we did not employ any filtering to refine localizations along the actin fiber as previously used in single molecule imaging^19^. Performing RCC, based on gold nanorods deposited onto coverslips, did not significantly reveal any drift during acquisition or improve the imaging resolution of our microscope (Fig. 3e and Supplementary Figs. 12 and 13).

**Figure 3:**
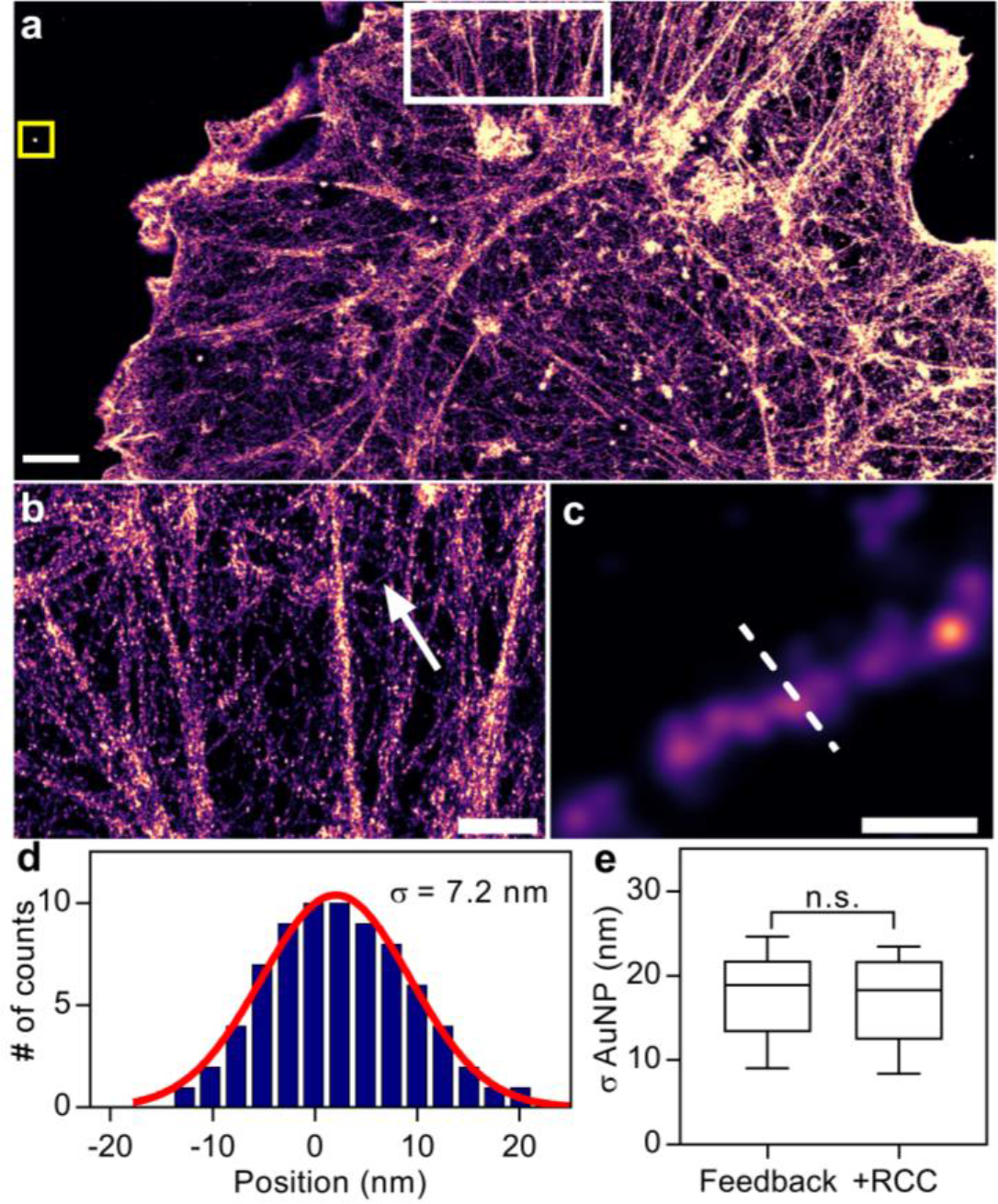
Feedback SMLM achieves ultrahigh resolution of cellular structures. **a**, Raw image of DNA-PAINT phalloidin in a COS-7 cell. Scale bar = 2 μm. **b**, Zoomed region of F-actin structures highlighted (white square) in (a). Scale bar = 1 μm. **c**, Zoomed region of a single actin filament indicated by the arrow in (b). Scale bar = 50 nm. **d**, Cross-sectional profile of an F-actin filament in (c). Red line, Gaussian fit with sigma of 7.2 nm. **e**, Size distribution of gold nanoparticles that were embedded in the sample (yellow square in (a)). Data are mean and standard deviations of n = 10 nanoparticles; n.s. not significant (P > 0.05, t-test assuming equal variance). Post-acquisition RCC did not improve the resolution or reduce drift.

An unsolved question in T cell biology is how antigenic peptides bound to major histocompatibility complex (pMHC) molecules initiates T cell receptor (TCR) signaling. It is generally assumed that TCR triggering requires the exclusion of phosphatases such as CD45^20^. However, so far it has not been possible to measure the separation between signaling TCR-CD3 complexes and CD45. Here, we activated Jurkat-ILA T cells on a supported lipid bilayer with antigenic pMHC-I molecules^21^ (Supplementary Fig. 14) and performed multi-channel acquisitions of CD45 and phosphorylated CD3ζ (pCD3ζ) with Feedback SMLM (Fig. 4a). As above, even unprocessed images revealed that CD45 was spatially excluded from the pCD3ζ nanoclusters (Fig. 4b). To measure the spatial separation, we performed a nearest neighbor distance (NND) analysis between pCD3ζ and CD45 in both directions and found a median distance of 19.6 nm (pCD3ζ to CD45) and 15.7 nm (CD45 to pCD3ζ) (Fig. 4c and Supplementary Fig. 15). NND on selected regions of high density of nanoclusters (n = 40 regions, 10 per cell) shows comparable distributions (Supplementary Figs. 16 and 17). More importantly, the NND analysis of Feedback SMLM data allowed us to plot the histogram of NND for 1-20 nm, the scale on which productive interactions between the phosphatase and its substrate can occur. We note that such distance measurements cannot be conducted with alternative approaches such as FRET or methods that rely on post-acquisition drift corrections using DNA origami structures or gold nanorods^22^ (Supplementary Figs. 18 and 19). In addition, the data quality of Feedback SMLM is unaffected by the inclusion of a supported lipid bilayer in the sample (Supplementary Fig. 20), which highlights the flexibility for sample preparation and SMLM data acquisition.

**Figure 4:**
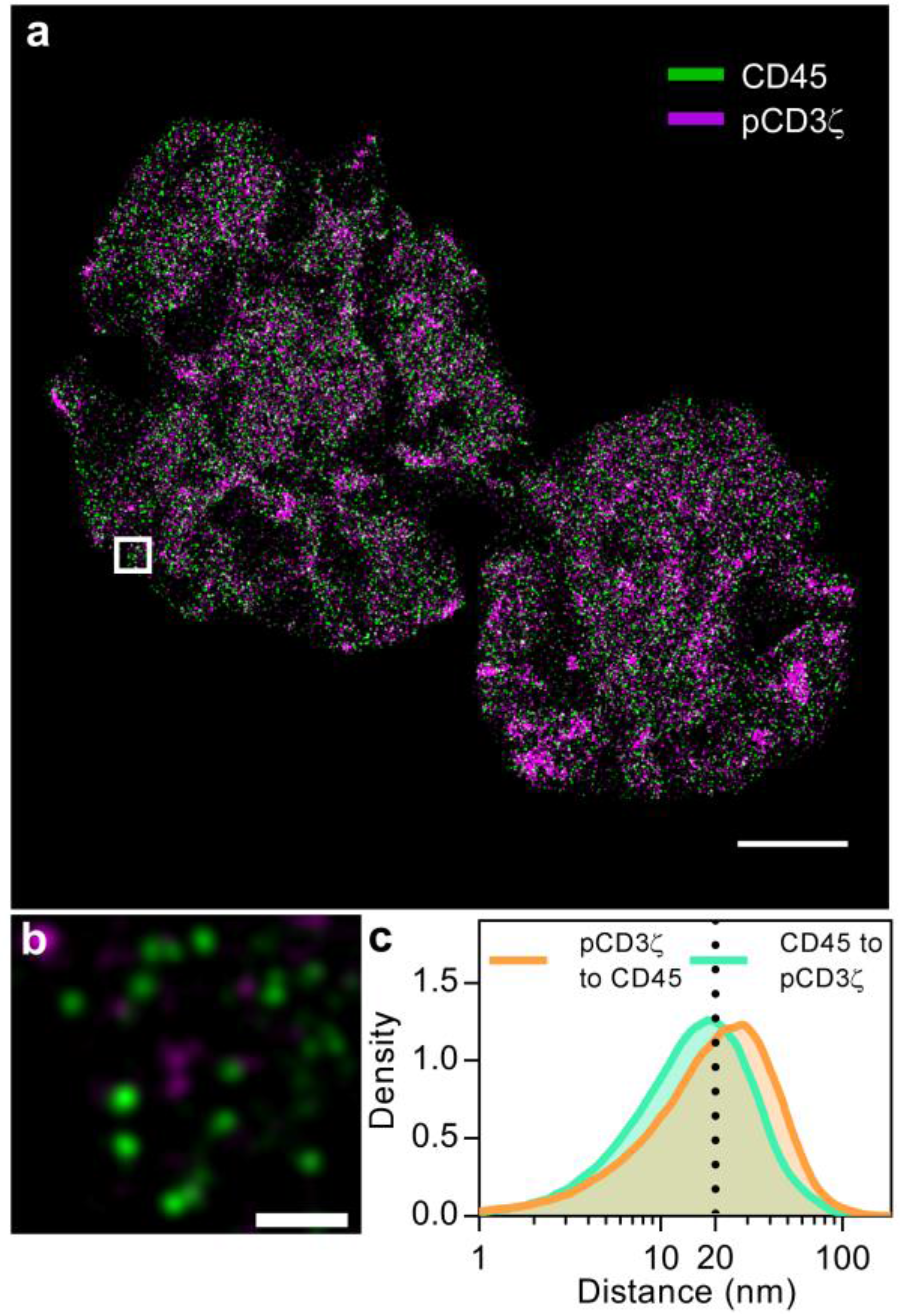
Distance measurements of interactions on biologically relevant scales. **a**, DNA-PAINT image of CD45 (green) and phosphorylated CD3ζ (pCD3ζ, magenta) in a Jurkat-ILA cell stimulated with a supported lipid bilayer presenting pMHC molecules and the adhesion protein ICAM1. Scale bar = 4 μm. **b**, Zoomed region of the highlighted area (white box) in (a) shows pCD3ζ nanoclusters were spatially separated from CD45. Scale bar = 200 nm. **c**, Nearest neighbour distance (NDD) for pCD3ζ to CD45 (orange line) and for CD45 to pCD3ζ (green line) with medians of 19.6 nm and 15.7 nm, respectively. The dotted line indicates 20 nm.

There has been debate in the field on whether nanometer and sub-nanometer localization accuracy can be practically achieved with SMLM^23^ given issues such as limited photon budget and unknown dipole orientation^24^. To successfully resolve structures such as DNA origami or the nuclear pore complex^25^, with comparable precision, assumptions are required to enable post-acquisition processing and averaging^26^. This prevents analysing structures of unknown shape or diverse structures with an unknown number of subunits. With Feedback SMLM such post-acquisition processing steps were not required, and yet structures such as individual F-actin filaments and molecular separation distances < 20 nm could be resolved. Thus, Feedback SMLM could become the tool of choice for biologists to quantify molecular associations and interactions on the nanometer scale.

## Acknowledgments

We thank Pierrette Michaux at the ANFF for the nanohole array. T. Boecking, M. Catarino and J. Goyette for assistance with the manuscript preparation.

## Funding

This work was supported by the Australia Research Council (CE140100011 to K.G.) and National Health and Medical Research Council of Australia (APP1059278 to K.G.).

## Author contributions

S.C., J.B. and K.G. wrote the paper. S.C., J.B., P.N. and K.G. conceived of the project. S.C. and J.B. built the Feedback SMLM. S.C, J.B and K.F. analyzed the experimental data. S.C., J.B., M.G. and J.H. performed the experimental work. J.H and H.G. developed the DNA-PAINT antibodies.

## Competing interests

The authors have no competing interests.

## Data and materials availability

Data and materials are available on request to the corresponding author.

## Code availability

Available on request to the corresponding author

## Methods

### Buffers

Imaging buffer A: Phosphate-buffered saline solution (PBS) with 10 mM magnesium. Cytoskeleton buffer B: 10 mM of MES, 150 mM of NaCl, 5 mM of EDTA, 5 mM of glucose and 5 mM of MgCl_2_. Buffer C: buffer B with 0.3% glutaraldehyde and 0.25% Triton x100. Buffer D: buffer B with 2% glutaraldehyde. Buffer E: Freshly prepared 0.1% NaBH4 in PBS. Buffer F: 10mM Tris - HCL, 100 mM NaCl, 0.05% Tween 20, pH 7.5. Buffer G: 5 mM Tris -HCL, 10 mM MgCl2, 1mM EDTA, 0.05% Tween 20, pH 8. Buffer H: PBS, 500 mM NaCl, pH 7.2.

### Preparation of glass coverslips

All coverslips were cleaned by sonication in ethanol for 15 min, followed by sonication in Milli-Q water for 15 min and plasma cleaning for 3 min, unless stated otherwise.

### Buffer exchange chamber

To perform buffer exchanges, we modified the lid of an 8-well chamber to accommodate an inlet and outlet port. Silicon tubing connected to syringes were inserted into the well via right angle brackets. Buffer present in the chamber was first removed via the outlet and the sample was washed before adding new buffer.

### DNA origami rulers on glass coverslips

A single well of an 8-well chamber (ibidi 80841) was attached to a clean coverslip and washed with 500 μl of PBS. The well was incubated with 200 μl of BSA-biotin solution (1 mg/ml in PBS) for 5 min. Excess BSA-biotin was removed by washing with 500 μl of PBS. The surface was incubated with 200 μl of neutravidin (1 mg/ml in PBS) for 5 min and washed with buffer A. Biotin-coated polystyrene beads (Spherotech, TP-30-5) (40 μg/ml) were incubated for 1 h and the excess beads were removed. The well was incubated with the DNA-origami ruler (GATTA-PAINT, HiRes 20R) diluted 40 times in buffer A to get ~100 rulers per field-of-view. Excess DNA origamis were removed by washing with buffer A. The imaging strand was a 9-bp complementary target strand with Atto 655, with a concentration of 5nM in buffer A.

### Cell culture

COS-7 cells were cultured in DMEM (Life Technologies, 11885-084) supplemented with 10% FBS, 1% penicillin-streptomycin. Jurkat-ILA1 T cells were cultured in RPMI 1640 (Life Technologies, 21870-076) supplemented with 10% FBS, 2 mM L-glutamine, 1 mM Penicillin and 1 mM streptomycin (all from Life Technologies). Characterization of the Jurkat-ILA1 T cells was performed as previously described^21^.

### Actin imaging

A coverslip with gold nanorods (HESTZIG, 600-30 AuF) was cleaned and a well chamber (ibidi 80841) was attached to the surface. The surface was coated with PLL-PEG-biotin (10 μg/ml in PBS, 20 min) to prevent non-specific binding, followed by streptavidin (0.09 μM). COS-7 cells were added (10,000-20,000 cells per well) and incubated for 24 h at 37 °C. Biotin-coated 3 μm polystyrene beads (Spherotech, TP-30-5) were mixed with serum-free DMEM (Life Technologies) and deposited onto the surface for 1 h at 37 °C and washed to remove excess beads. Cells were fixed and permeabilized with 300ul of buffer C for 1-2 min, followed by 600ul of buffer D for 10min. Cells were treated with 600 μl of buffer E for 7 min and washed with PBS. The surface was passivated using biotin in 5% BSA (1 μM) for 1 h and washed with PBS. Cells were incubated overnight at 4 °C with 300 μl of Biotin-XX phalloidin (Thermofisher, B7474) (0.5 μM) in 5% BSA and then washed with buffer F. Buffer F with streptavidin (3.45 μM) was added to the cells at room temperature for 30 min and washed with PBS. Buffer D was added for 10 min at room temperature. We then washed with PBS, buffer F and buffer G. Biotinylated DNA docking strands (2 μM) in buffer G were incubated for 30-60 min at room temperature and washed with buffer G. Imaging strands (500 pM) in buffer H was added and the sample imaged.

### Bilayer Preparation

Glass coverslips were cleaned with 1M KOH for 10 min, rinsed with MilliQ water, placed in 100% ethanol for 20 min and plasma cleaned for 5 min. 8 well silicone chambers (ibidi, 80841) were then attached to the cleaned coverslip. A liposome solution of 1mg/ml with a lipid ratio of 96.5% DOPC (1,2-dioleoyl-*sn*-glycero-3-phosphocholine), 2% DGS-NTA(Ni) (1,2-dioleoyl-*sn*-glycero-3-[(N-(5-amino-1-carboxypentyl)iminodiacetic acid)succinyl] (nickel salt)), 1% Biotinyl-Cap-PE (1,2-dioleoyl-*sn*-glycero-3-phosphoethanolamine-N-(cap biotinyl) (sodium salt)), and 0.5% PEG5000-PE (1,2-distearoyl-*sn*-glycero-3-phosphoethanolamine-N-[methoxy(polyethylene glycol)-5000] (ammonium salt) (mol%; all available from Avanti Polar Lipids (DOPC, 850375C), (DGS-NTA(Ni), 790404C), (Biotinyl-Cap-PE, 870273C), (PEG5000-PE, 880220C) was created by vesicle extrusion, as described in detail elsewhere ^27^. The lipid solution was added to the wells at a 1:5 ratio with MilliQ water along with 10mM of CaCl_2_ for 15 min and washed repeatedly with PBS. Followed by 0.5mM EDTA in MilliQ water to remove the excess CaCl_2_. The well was then washed with PBS and incubated with 1mM of NiCl_2_ to recharge the NTA groups. Disruption of the lipid bilayer was avoided by maintaining 100-150 μl of PBS in the wells.

### Bilayer decoration for DNA origami rulers

To decorate the bilayer with fiducials and DNA origami rulers, 200 μl of neutravidin (1mg/ml in PBS) was incubated in the well for 5 min and washed with buffer A. Biotinylated polystyrene beads (Spherotech, TP-30-5) were then added at 40 μg/ml for 30 min and then washed with buffer A to remove excess beads. The well was then incubated with biotinylated nanorods (NanoPartz, C12-25-650-TB-DIH-50) at 1:20000 ratio and DNA-origami rulers (GATTA-PAINT HiRes 20R) diluted 40 times in buffer A and washed with buffer A to remove any unbound fiducials and rulers. The well was fixed with 4% PFA for 15 min and washed with buffer A. The imaging strand which consists of Atto655 on a 9-bp complementary target strand was added to the chamber at a concentration of 5nM in buffer A.

### Bilayer decoration for T cell activation

To decorate the bilayer with fiducials and proteins, 100 μg/ml of streptavidin (Life Technologies, SNN1001) was incubated for 10 min and washed with PBS. Biotinylated polystyrene beads (SpheroTech, TP-30-5) were then added at 40 μg/ml ratio in PBS for 20 min and washed with PBS to remove excess beads. Biotinylated pMHC 3G (500 ng/ml), His-tagged ICAM-1 (200 ng/ml,) and biotinylated nanorods (NanoPartz, C12-25-650-TB-DIH-50) at a 1:20000 ratio were combined with 5% BSA in PBS. Biotinylated and His-tagged proteins and biotinylated nanorods were then added to the well for 30 min. Finally, the surfaces were washed with PBS to remove unbound fiducials or proteins.

### T cell activation on bilayer and immunostaining

The wells were washed with RPMI culture media and warmed to 37 °C for 30 min. Jurkat-ILA cells were added to the bilayer at a density of 250,000 cells in 50 μl for 4 min at 37 °C and fixed using 4% PFA in PBS for 20 min at 37°C. For immunostaining, cells were permeabilized with Triton X-100 (Sigma-Aldrich, T8787) at 0.1% for 5 min at room temperature and washed with PBS. The cells were blocked with 5% BSA in PBS for 1 h. Cells were labelled with primary antibodies against pCD3ζ (pY142) (BD Pharmingen, 558402), and CD45 (Abcam, ab10559) for 30 min, then washed with PBS. Primary antibodies were previously conjugated with DNA strands (Table S1). Cells were then labelled with a secondary antibody against pCD3ζ (Jackson Immunoresearch, 115-547-003) for 30 min at room temperature, and then washed with PBS. The sample was fixed using 4% PFA in PBS for 15 min and then washed with buffer A. DNA target strands conjugated in buffer A were added to the sample at a 5 nM concentration. CD45 and pCD3ζ docking strands are all complementary to the individual docking strands on the primary labelled antibodies. Imaging was performed sequentially after performing buffer exchanges.

### Optical setup

Laser lines of 405 nm (Vortran, Stradus 405-100), 488 nm (Vortran, Stradus 488-150) and 637 nm (Vortran, Stradus 637-180) were passed through clean-up filters (Semrock, 405 nm: LD01-405/10-12.5; 488 nm: LL01-488-12.5, 640 nm: LD01-640/8-12.5), combined into a single path using dichroic mirrors (Chroma; ZT442rdc and ZT594rdc) and fiber-coupled (Thorlabs, P3-405BPM-FC-2). A pair of achromatic doublet lenses (f = 30 mm and f = 300 mm) were used to expand the lasers. The lasers were focused onto the back aperture of a 100× 1.49 NA TIRF objective (Nikon, CFI Apochromat) using an achromatic lens (f = 200 mm). TIRF illumination was achieved by displacing the laser beams towards the periphery of the objective. The displacement was performed by moving the focusing lens with a mirror assembled on a translation stage (Newport, M-423-MIC). Lasers were delivered to the objective using a dichroic beamsplitter (Chroma, ZT488/640rpc), which reflected all lasers (and infrared LED) but allowed for transmission of the fluorescence. The sample was mounted on a nano-positioning stage with 0.1 nm step size in *x/y* and 0.4 nm in the z-axis (Mad City Labs, LP50-200), integrated on an inverted microscope body (Mad City Labs, RM21). The optical setup was built on an actively stabilized optical table (Newport, M-ST-46-8) and tuned for the payload distribution. Table vibrations were monitored in real-time (Newport, ST-300).

Sample fluorescence was filtered using an emission filter (Semrock, Em01-R405/488/635-25). A lens (f = 400 mm) focused the fluorescence onto an EMCCD camera (Andor, iXon 897 Ultra) via a piezoelectric mirror (Thorlabs, Polaris-K1S3P). The EMCCD camera was water cooled (Koolance, EX2-1055) to avoid fan vibrations and mounted in a custom-made steel holder. The EMCCD camera was set to a temperature of −95°C to minimize camera pixel noise. The final magnification of the EMCCD camera resulted in a 40 μm × 40 μm field-of-view (individual pixel size = 80 × 80 nm). A white LED (Mightex, BLS-LCS-4000-03-22) illuminated a pinhole and was combined with the fluorescence path using a 0.5% reflective mirror (Thorlabs, WG11010-A). The spot on the EMCCD camera created by the LED was fitted with a 2D Gaussian and deviations from the original position were applied to the piezo-electric mirror to correct for drift in the fluorescence path. To avoid disturbance to the fluorescence, the LED spot was positioned on the periphery of the EMCCD camera. Image deviation was calculated as (*dx*^2^+*dy*^2^)^0.5^ for each frame, where *dx* and *dy* represent the deviations registered from the set position.

To perform sample stabilization, an infrared LED (Mightex LCS-0850-02-22) was filtered using a bandpass filter (Semrock, FF01-842/56-25) and directed to the sample via a condenser (Thorlabs, ACL25416U-B). The infrared light was removed from the fluorescence path using the laser dichroic, separated from the lasers using a dichroic beamsplitter (Semrock, FF801-Di02-25x36), filtered using a bandpass filter (Semrock, FF01-842/56-25) and focused (f = 200 mm) onto a CMOS camera (Allied Vision Manta G-235). The final magnification of the CMOS camera resulted in a 112 μm × 70 μm field-of-view (individual pixel size = 58.6 nm), ~5-fold larger than the EMCCD camera. The non-coherent light source did not generate interference speckles^28^ and produced a high contrast diffraction pattern on the CMOS camera. The focus of the CMOS camera is offset by 3 μm with respect to the fluorescence camera.

The data acquisition and instrument control were performed using an Xbox One controller (Microsoft) integrated into 64-bit LabVIEW (National Instruments).

### Environmental control

An environmental control box was installed around the optical setup to control temperature fluctuations (Supplementary Fig. 21). A series of Peltiers (Thorlabs, TEC3-6) with temperature transducers (Thorlabs, AD590) were regulated individually (Thorlabs, TED200C). Temperature and humidity were registered independently from the temperature control units (Thorlabs, TSP01). The temperature and humidity show a standard deviation of 0.02 °C and the 0.88%, respectively. Prior to imaging, we allowed the samples to acclimatize for 15-30 min.

### Active stabilization

A clean glass coverslip was coated with PLL-PEG-Biotin (10 μg/ml in PBS) to prevent non-specific binding, followed by streptavidin (0.09 μm). Biotin-coated polystyrene beads (Spherotech, TP-30-5) (40 μg/ml) were incubated for 1 h and the excess beads were removed. Surface passivation was performed with biotin (1 μm) and then washed with PBS.

The software can automatically identify polystyrene beads within the field-of-view (FoV). The center of the particle was identified (within ~1 pixel) by a background-corrected center of mass (COM) algorithm^14^ and produced a square region-of-interest (ROI) of 150 × 150 pixels around each bead. This initial calculation was not critical as it served as a visual aid that allowed the software to identify beads and register ROIs. CMOS technology allows for an increased frame rate by selectively recording a ROI. To correct for drift (and monitor residual drift), the ROIs of two beads were recorded. One bead was referred to as the lock bead and a second bead was referred to as the reference bead. The software calculated the position of both beads, however only acted upon drift registered by lock bead. The reference bead served as an internal control, or an out-of-loop reference, to monitor drift (Fig 1C-D and Supplementary Fig. 2). The average drift over a period of 10 frames was subtracted to the stage position. Applying an equal but opposite drift amplitude suppressed drift movement. The stage was moved at a frequency of 10-15 Hz to adequately correct for sample drift. Real-time 3D drift correction was performed using a camera-based particle tracking software implemented in LabVIEW. We employed real-time data processing in a graphics-processing unit (Nvidia, GTX1070), which has been shown to achieve 3D particle tracking of fixed beads with a precision of 0.1 nm at kHz rates and down to 0.01 nm at 10Hz rates in all dimensions^15^. The lateral positions of the bead are determined by correlating linear bead intensity profiles with their mirror profiles^13^. The axial position was determined by comparing the radial intensity distribution to a pre-recorded look-up table (LUT). The LUT that served as an axial calibration was acquired by performing a z-stack over a 4 μm depth by stepping the nanopositioning stage in 100 nm increments^13^. The software allowed for image streaming together with the lateral/axial position determination to be carried out in parallel^15^.

### Standard SMLM microscope

Standard SMLM imaging was performed on a commercial Zeiss Elyra SMLM microscope. TIRF illumination from a 640 nm laser diode was focused on DNA origami structures via a 100× 1.49 NA Zeiss objective. Axial drift correction was performed in real-time using the Zeiss focus-lock module (Zeiss, Definite Focus). Fluorescence was filtered and captured on an EMCCD camera (Andor, iXon 897 Ultra) with a pixel size of 97 nm. The temperature was stabilized using the Zeiss environmental control module (Zeiss, TempModule S).

### Image processing and single-molecule localization

Individual fluorescent images were recorded and saved in a 16-bit TIFF format. The images were automatically split into 4 GB folders and an ImageJ macro was used to convert the images into a stack format (.OME.TIFF). To localize molecules, the PSF of an individual molecule within each frame was fitted to a 2D Gaussian. Images were analyzed with the Picasso software^9^ or ImageJ (NIH) with the ThunderSTORM plugin^29^. Cross-sectional profiles were fitted with a Gaussian or Lorentzian, according to their distribution^30^.

### Redundant cross-correlation

Post-acquisition drift correction was performed using the redundant cross-correlation algorithm (RCC). Previously reported RCC performed on simulated data has shown a root-mean square error of 1.5 nm^16^. Temporal bins were set at 1000 frames. Performing successive rounds of RCC with sequentially smaller temporal bins did not show an improvement in the data.

### Nearest neighbor distance analysis

To evaluate the distribution of pCD3ζ and CD45, the cross nearest neighbor distance (pCD3ζ to CD45; and CD45 to pCD3ζ) was calculated using R package ‘spatstat’. The distance was log transformed (base 10) before the density was calculated and plotted. Cell boundaries were delimited to eliminate erroneous calculations from the background.

### EMCCD characterization

To reduce the impact of systematic errors in the microscope’s detection path we adopted a strategy that realigns molecular positions^19, 31^. We produced a regular pattern with stable emission by generating a ‘nanohole array’ (Supplementary Fig. 7). The nanohole array consists of a series of circular holes with a diameter of 100 nm nanofabricated on an aluminum coated coverslip in a 12×12 array. The holes were arranged in grid pattern with a vertical and lateral separation of 1 μm. The holes were then filled with red (Alexa647) and green (Alexa488) dyes at ~100 μM concentration. We registered the pre-determined emission pattern in both the green and red channels. The location of the points was determined with nanometer accuracy. The apparent variation between the green and red coordinates was due to the registration errors of the EMCCD, optical and chromatic aberrations. The local deviations of each green-red point were registered as a vector displacement. The pattern was moved in an *xy* raster pattern in an overlapping approach to characterize the pixels of the EMCCD camera.

## Supplementary Information

**Supplementary Figure 1.**
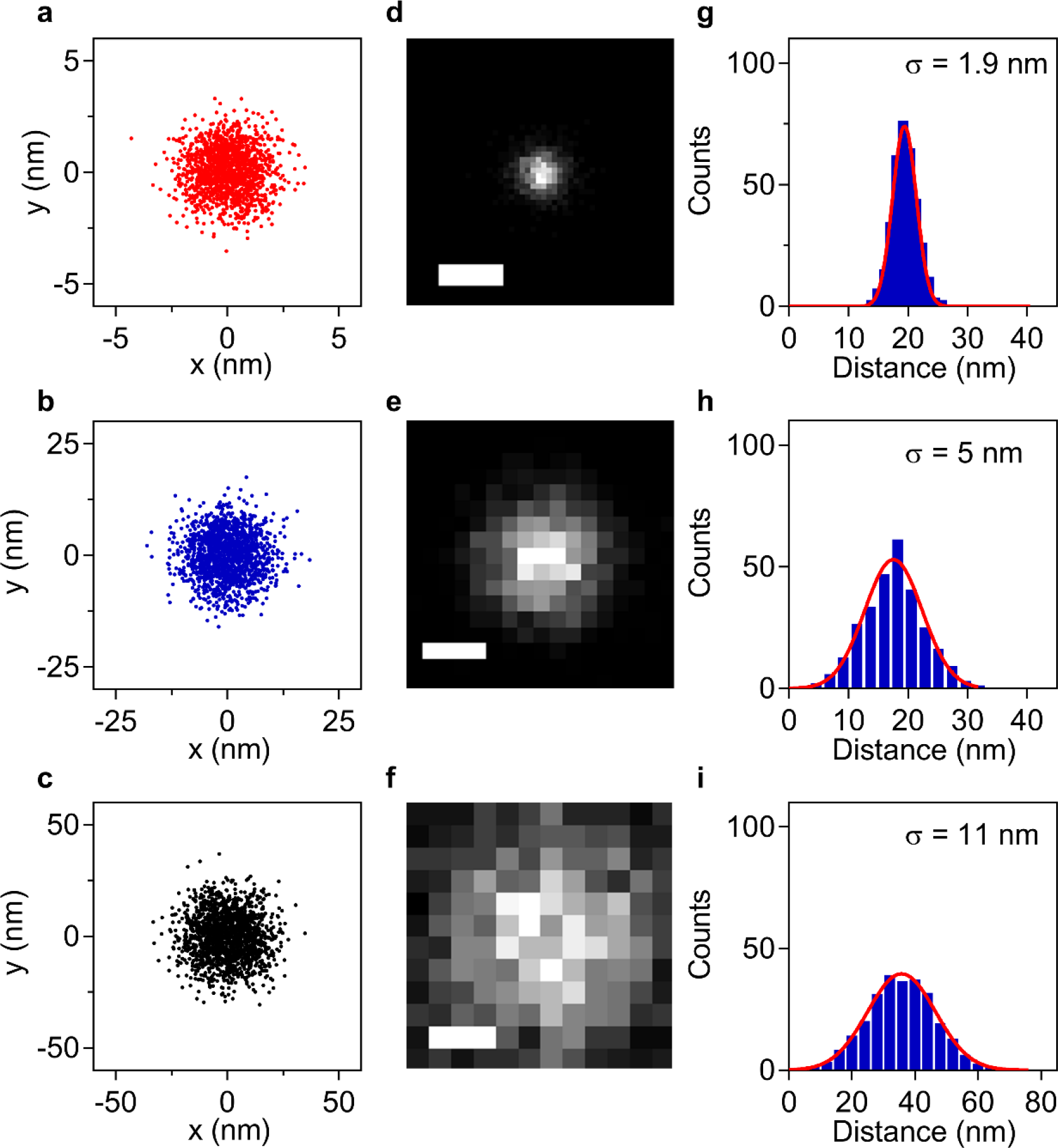
Degradation of localization precision as a function of drift. (**a-c**) Drift during acquisition was simulated by generating Gaussian noise with multiple standard deviations (SD) (N _Points_ = 2000). Simulated drift of 1 nm in (**a**), 5 nm in (**b**) and 10 nm in (**c**). (**d-f**) Simulated single molecule data with 1.5 nm localization precision was corrupted by the simulated drift. Scale bar = 10 nm. The magnification was adjusted to aid visualization. (**g-i**) Fitted cross-sectional line profiles.

**Supplementary Figure 2.**
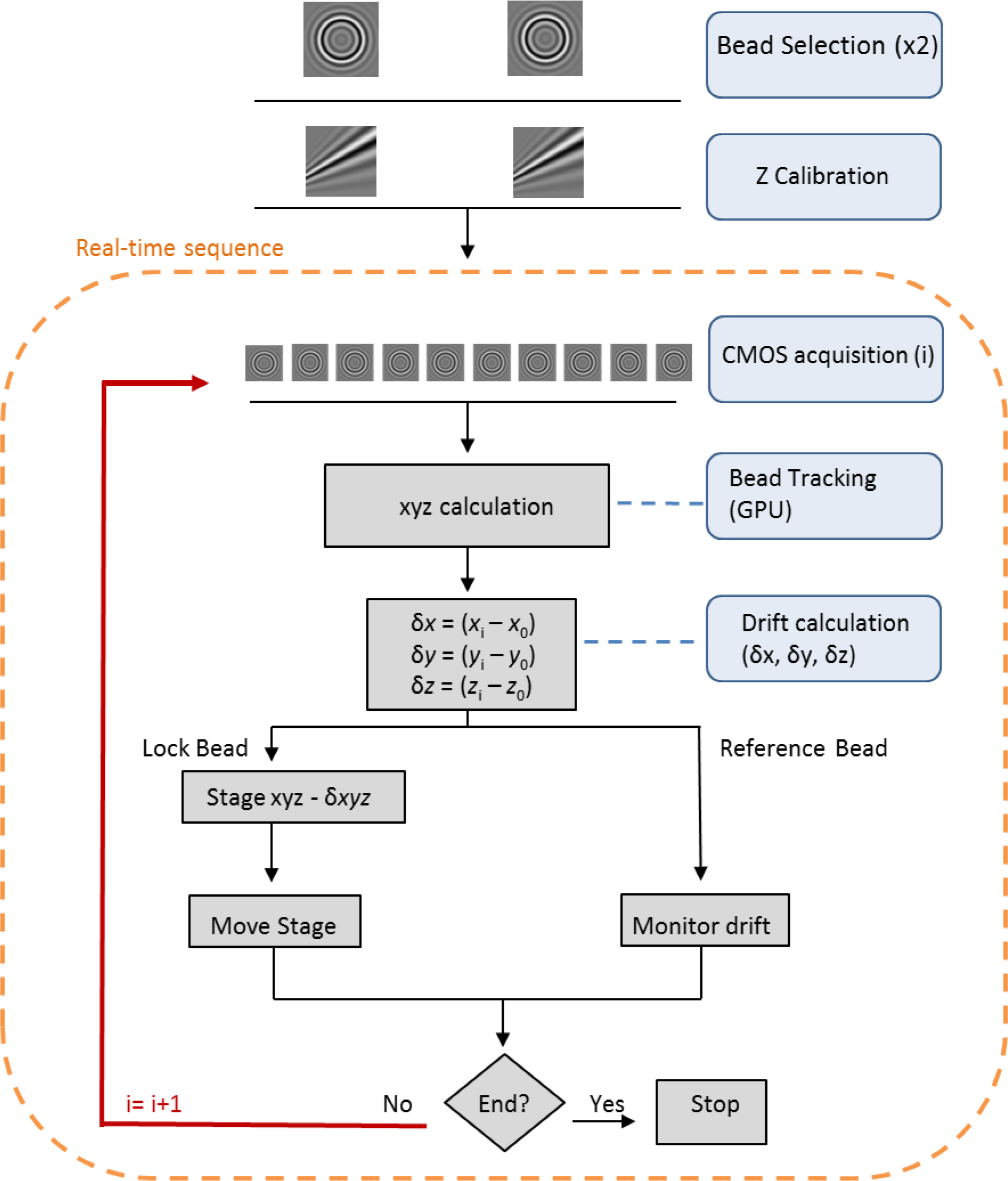
Active stabilization procedure. Real-time 3D active stabilization was performed and monitored by tracking two 3 μm polystyrene beads simultaneously (lock and reference bead). First, two beads were selected, and the axial calibration table acquired for both beads. The CMOS camera acquired a region of interest of the lock bead and determined the position using the GPU. The drift in each frame was calculated by subtracting the position of the lock bead from its original location. The stage was then moved to apply an equal but opposite displacement, therefore correcting for drift of the sample. The reference bead acted as an internal control to monitor sample stability.

**Supplementary Figure 3.**
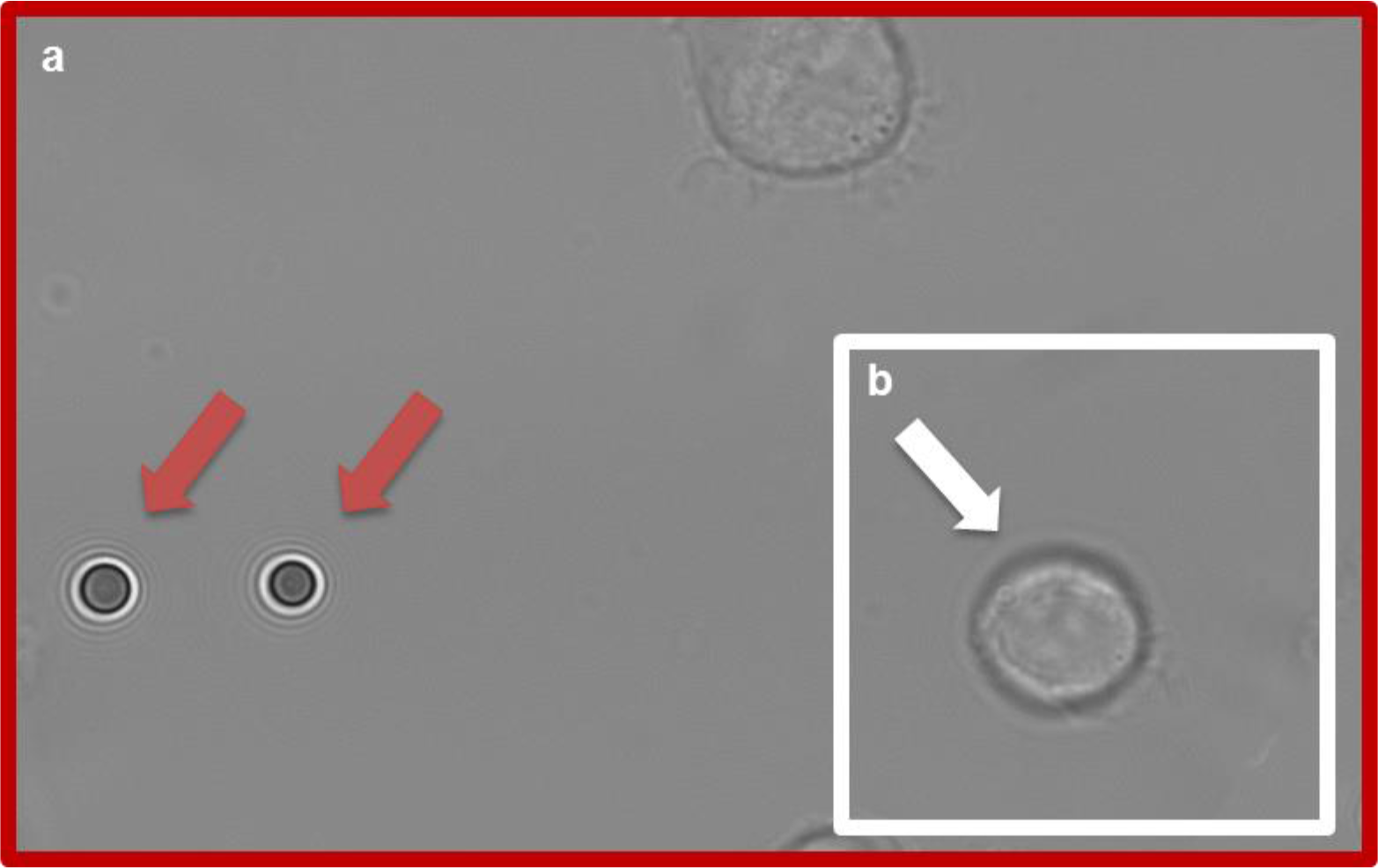
Fields-of-views of the Feedback SMLM. Non-fluorescent fiducials were used outside the field of view (FoV) of the cameras for SMLM (white square). This provided great flexibility in sample positioning and ensured that fiducials did not interfere with data acquisition. (**a**) An infrared LED illuminated the sample and the polystyrene beads on the coverslip (red arrows). The beads created diffraction rings which were imaged with a CMOS camera with a FoV of 112 μm × 70 μm (red square). (**b**) FoV of the fluorescent EMCCD camera was ~5 fold smaller (40 μm × 40 μm). The white arrow points to a T cell in the FoV of the SMLM camera.

**Supplementary Figure 4.**
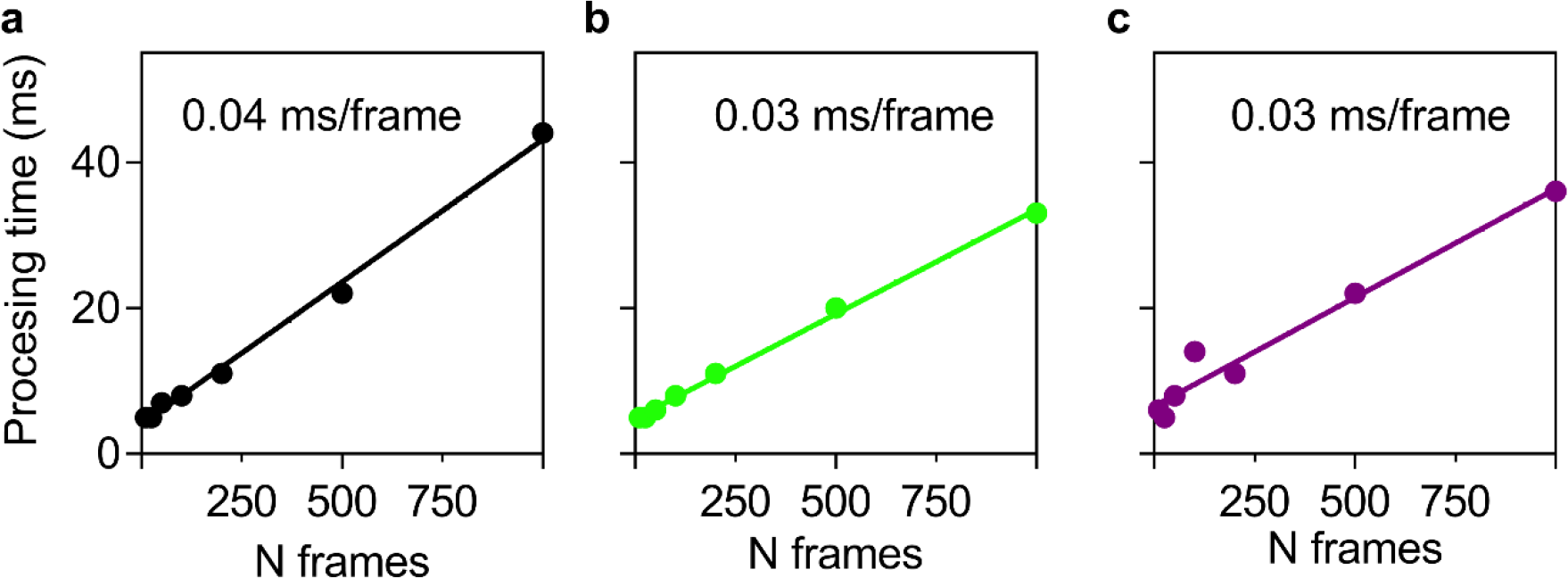
Speed of GPU calculation. The 3D position of a simulated bead was calculated for a varying number of frames. (**a**) Processing time without pixel noise for a stationary bead. (**b**) Processing time without pixel noise for a mobile bead. (**c**) Processing time with pixel noise for a mobile bead.

**Supplementary Figure 5.**
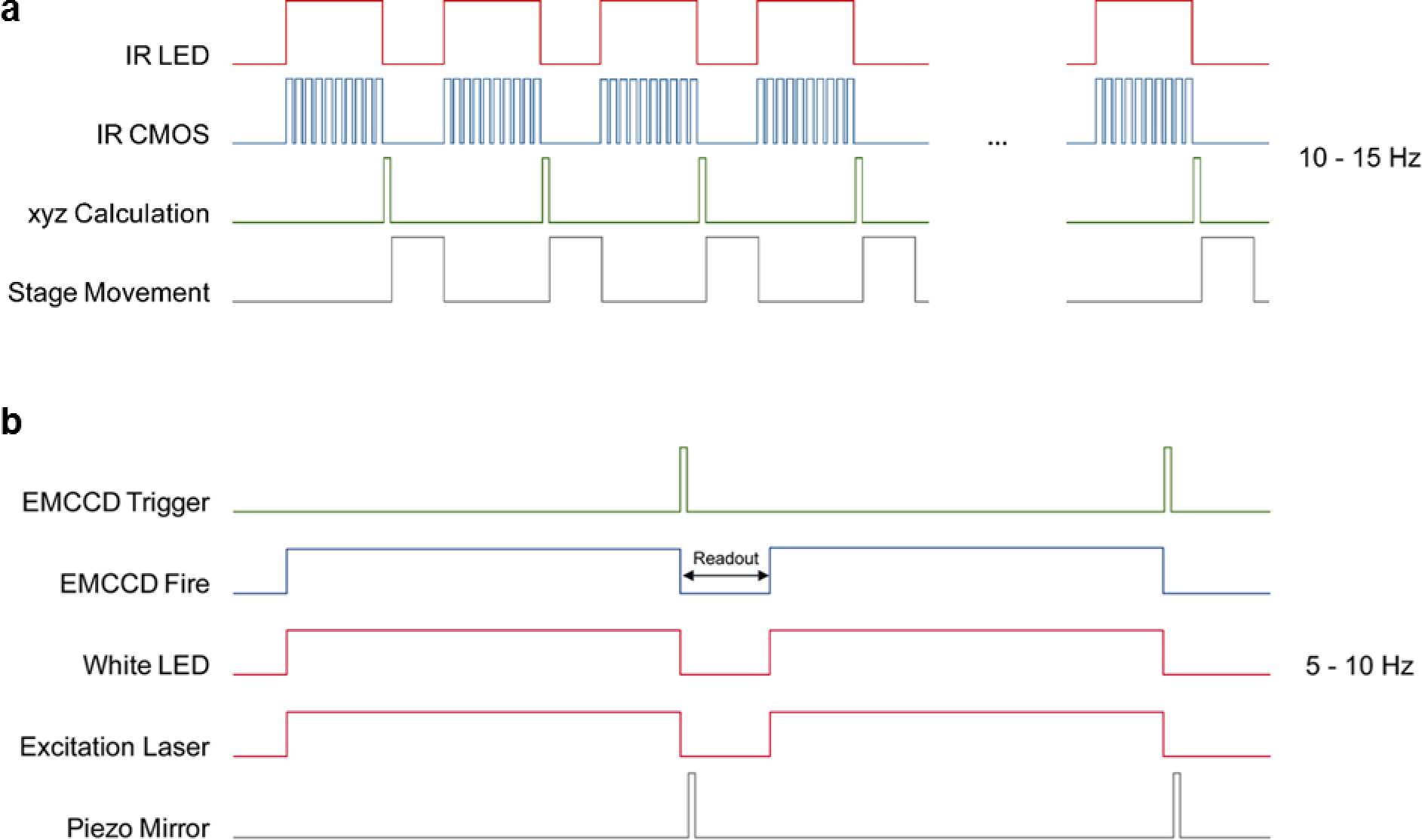
Acquisition diagrams. (**a**) Diagram of the timing of sample stabilization during acquisition. The stage movement was synchronized with the acquisition of the CMOS camera. (**b**) Diagram of the timing of the fluorescence acquisition. To enhance acquisition speed, the piezo electric mirror was moved during the readout of the EMCCD.

**Supplementary Figure 6.**
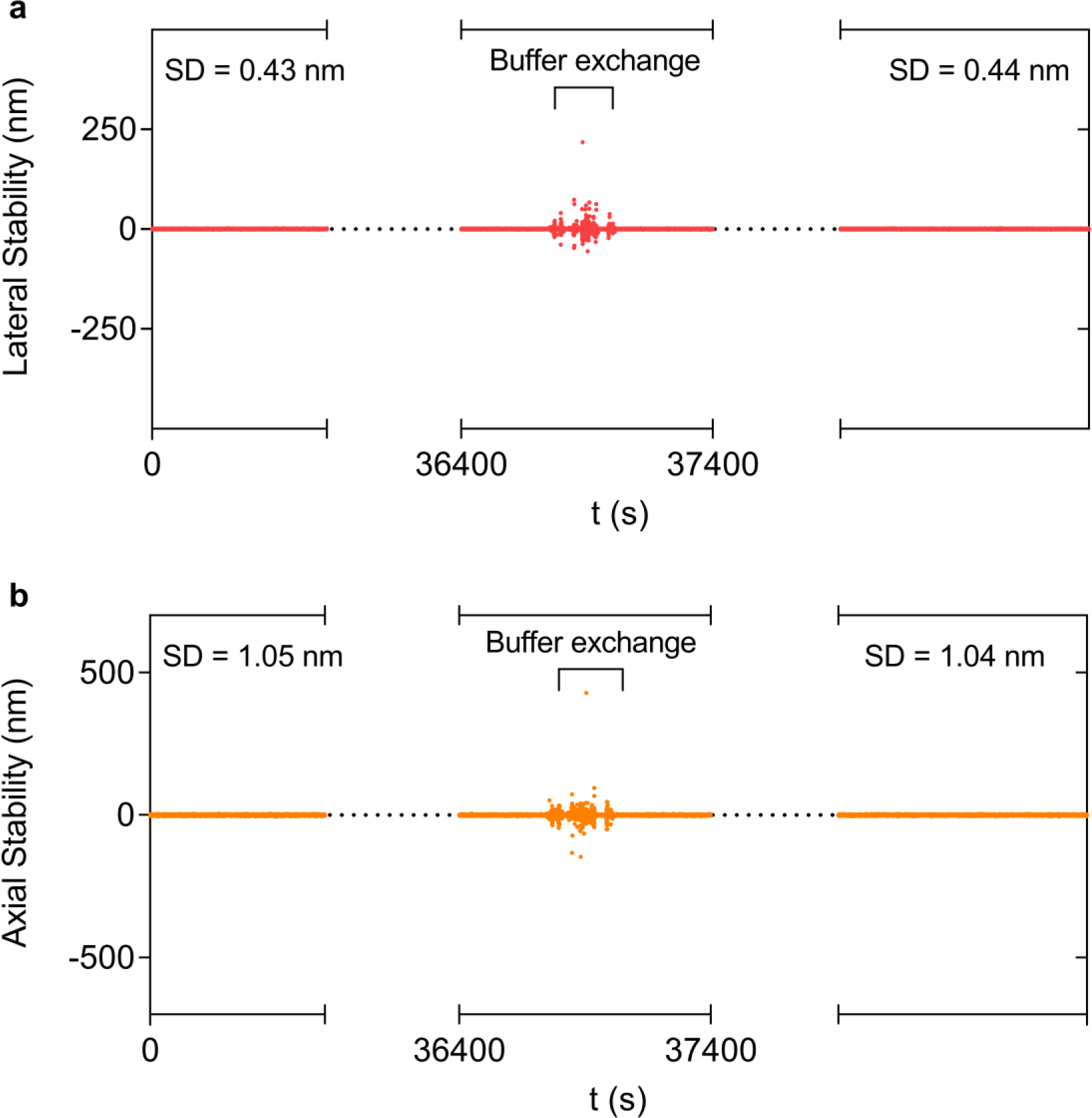
Feedback SMLM corrects stage position during buffer exchanges. Beads were attached to a lipid bilayer and incubated overnight. A buffer exchange was performed which induced movement of the sample. The Feedback SMLM autonomously repositions the sample and corrects for drift introduced. (**a**) Lateral stabilization. Pre- and post-buffer exchange shows a standard deviation of 0.43 nm and 0.44 nm. (**b**) Axial stability. Pre- and post-buffer exchange shows a standard deviation of 1.05 nm and 1.04 nm.

**Supplementary Figure 7.**
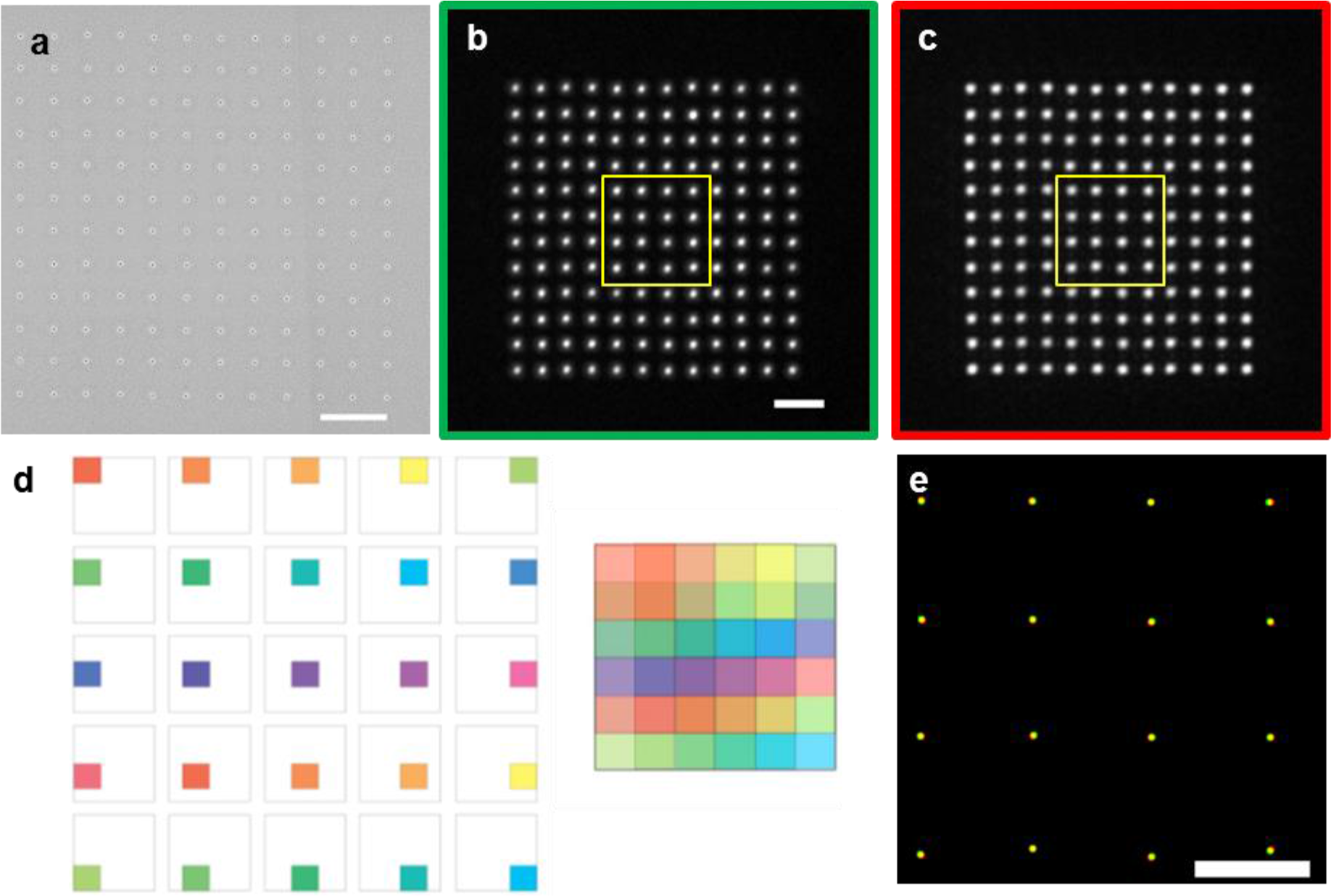
EMCCD camera characterization and correction. (**a**) Electron microscopy image of nanohole array used to test the EMCCD cameras. The nanohole array consists of sub-diffraction-sized circular holes in an aluminum layer on a glass coverslip arranged in a regular grid pattern and separated by 1 μm. Scale bar = 2 μm. (**b-c**) The nanohole array was filled with green (Alexa488, b) red (Alexa647, c) dyes. Scale bar = 2 μm. (**d**) Graphical illustration of the camera testing procedure. The nanohole array which occupied 1/9 of the EMCCD area was stepped in discrete increments. Each step was equal to half the length of the nanohole array in both the horizontal and vertical directions. Stepping the sections over the entire space generated a final overlapped mask in which each overlapping segment contributes equally. (**e**) Residual errors after applying the correction was reduced ~10-fold. Localizations from the yellow square highlighted in (b) and (c) after EMCCD correction. Scale bar = 1 μm.

**Supplementary Figure 8.**
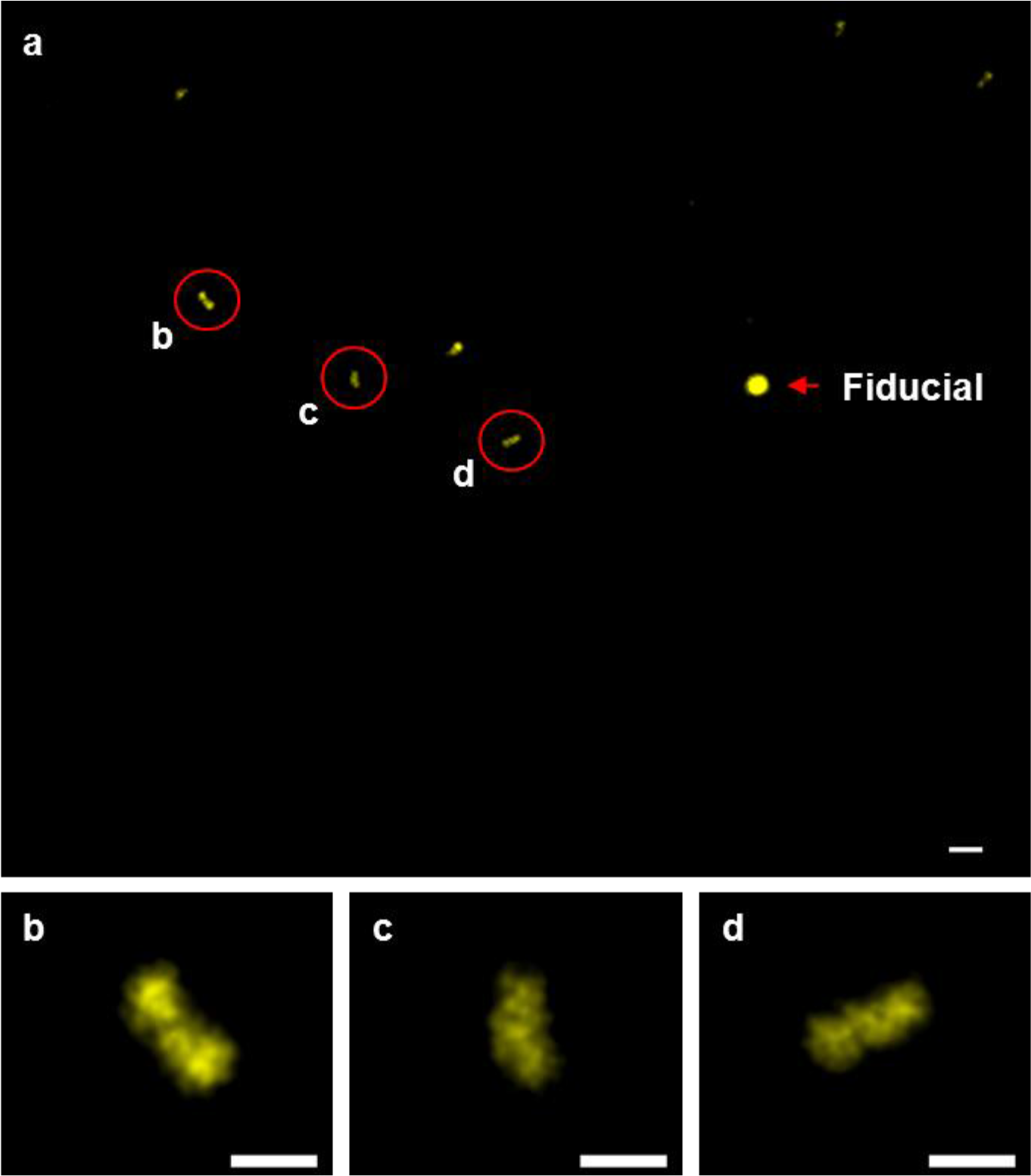
DNA-PAINT imaging of DNA origami structures with standard SMLM microscopy. (**a**) Images of DNA origami structures with standard SMLM microscopy after drift correction. Post-acquisition drift correction was performed using RCC and was essential to visualize the structures. Scale bar = 100 nm (**b-d**) Zoomed regions of individual origami rulers highlighted by the red circles in (A). Scale bars = 40 nm.

**Supplementary Figure 9.**
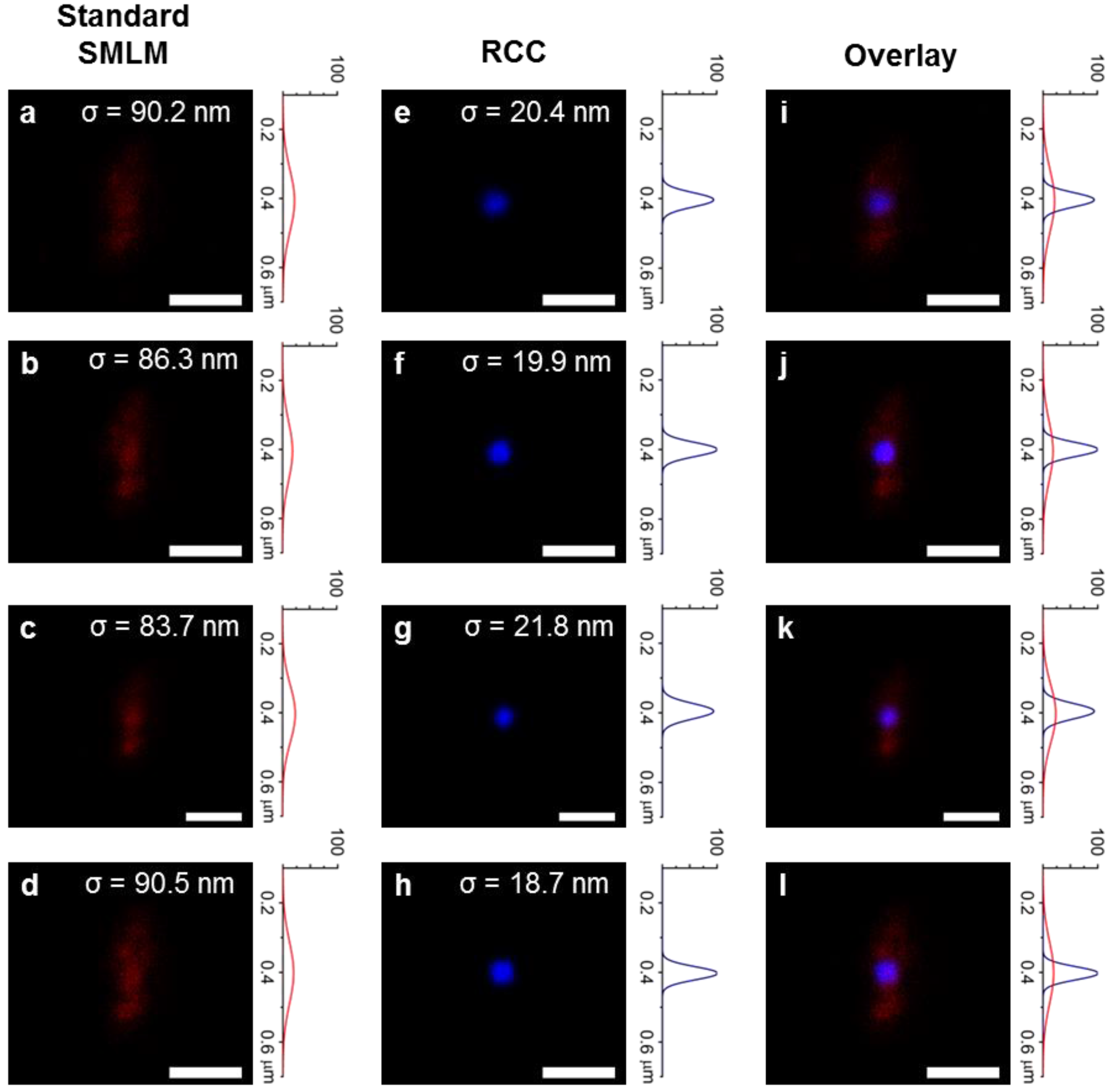
Influence of drift in standard SMLM microscopy. Gold nanorods were deposited on a glass coverslip to monitor drift. (**a-d**) Raw data from a standard SMLM acquisition without the active stabilization of Feedback SMLM. (**e-h**) Data from (a-d) was drift corrected using RCC. (**i-l**) Data overlaid from (a-d) and (e-h). The distribution of the gold nanorods were fitted to Gaussian curves (a-l). Scale bar = 200 nm.

**Supplementary Figure 10.**
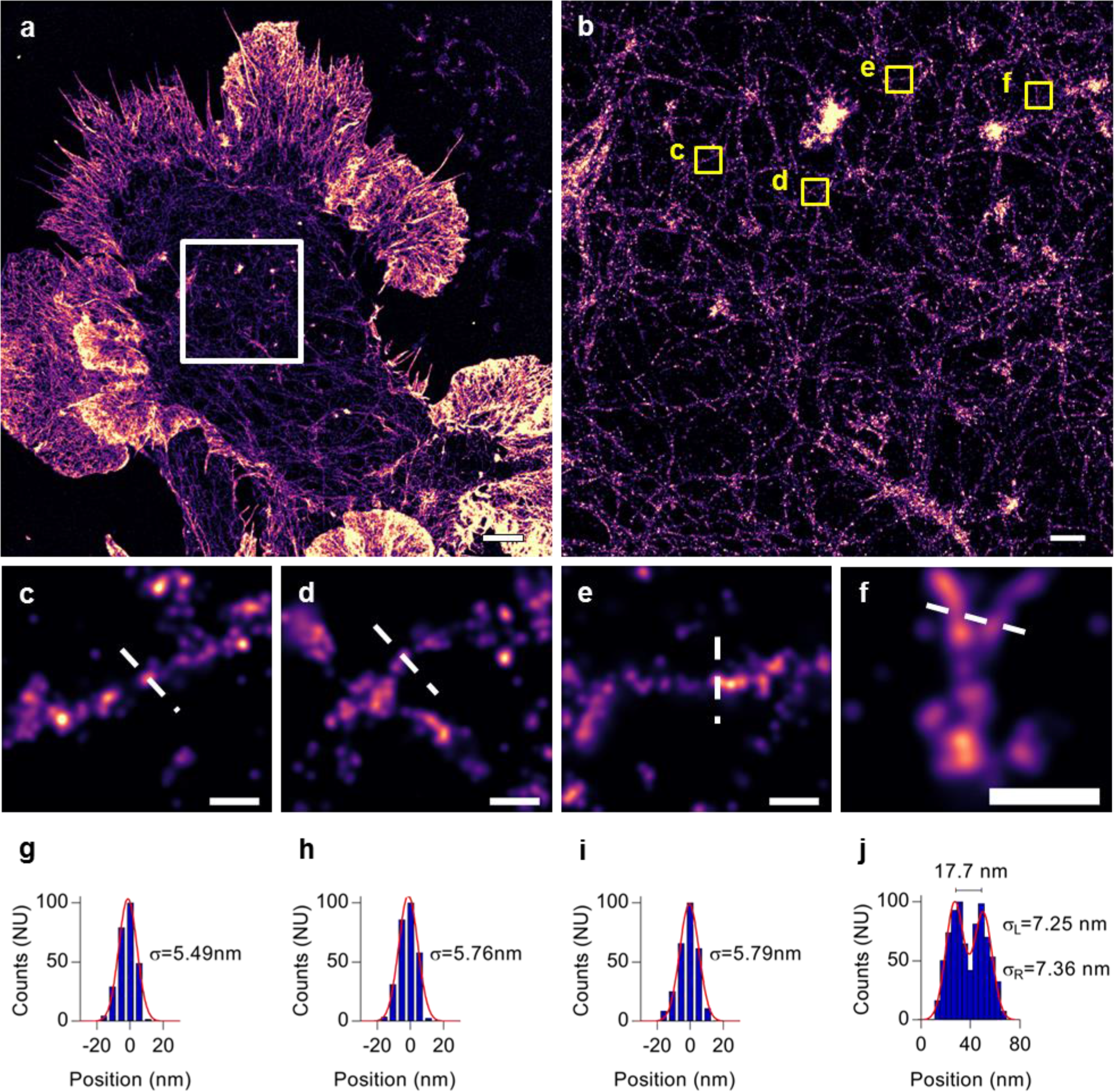
Raw data of F-actin in COS-7 cells obtained with Feedback SMLM. DNA-PAINT imaging of phalloidin-stained COS-7 cells with Feedback SMLM. Localisations were not filtered, grouped or altered post-acquisition. (**a**) Raw data from actively stabilized Feedback SMLM. Scale bar = 3 μm. (**b**) Zoomed region of the highlighted area (white square) in (a). Scale bar = 500 nm. (**c-f**) Image of selected, individual filaments (yellow squares) indicated in (b). Scale bars = 50 nm. (**g-j**) Cross-sectional Gaussian fit of actin filaments shown in (**c-f**).

**Supplementary Figure 11.**
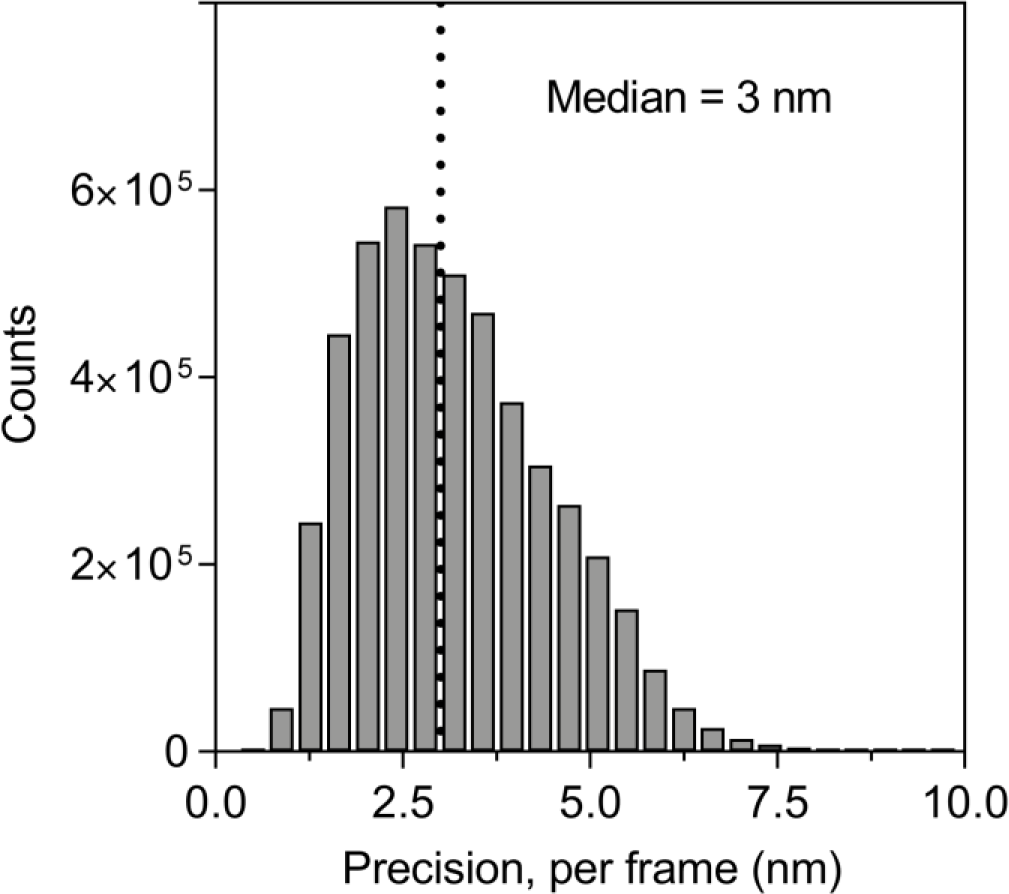
Histogram of localization precision per frame in F-actin for the DNA-PAINT images shown in Supplementary Figure 10. The median localization precision of the Atto655-conjugated imaging strand in a single frame was 3 nm (dotted line). Grouping across frames of fluorescent events, which belong to the same molecule would further improve the localization accuracy. Gold nanoparticles were excluded as well as localizations with excessive photon counts and uncharacteristic binding events. The number of localizations represented was ~5 million.

**Supplementary Figure 12.**
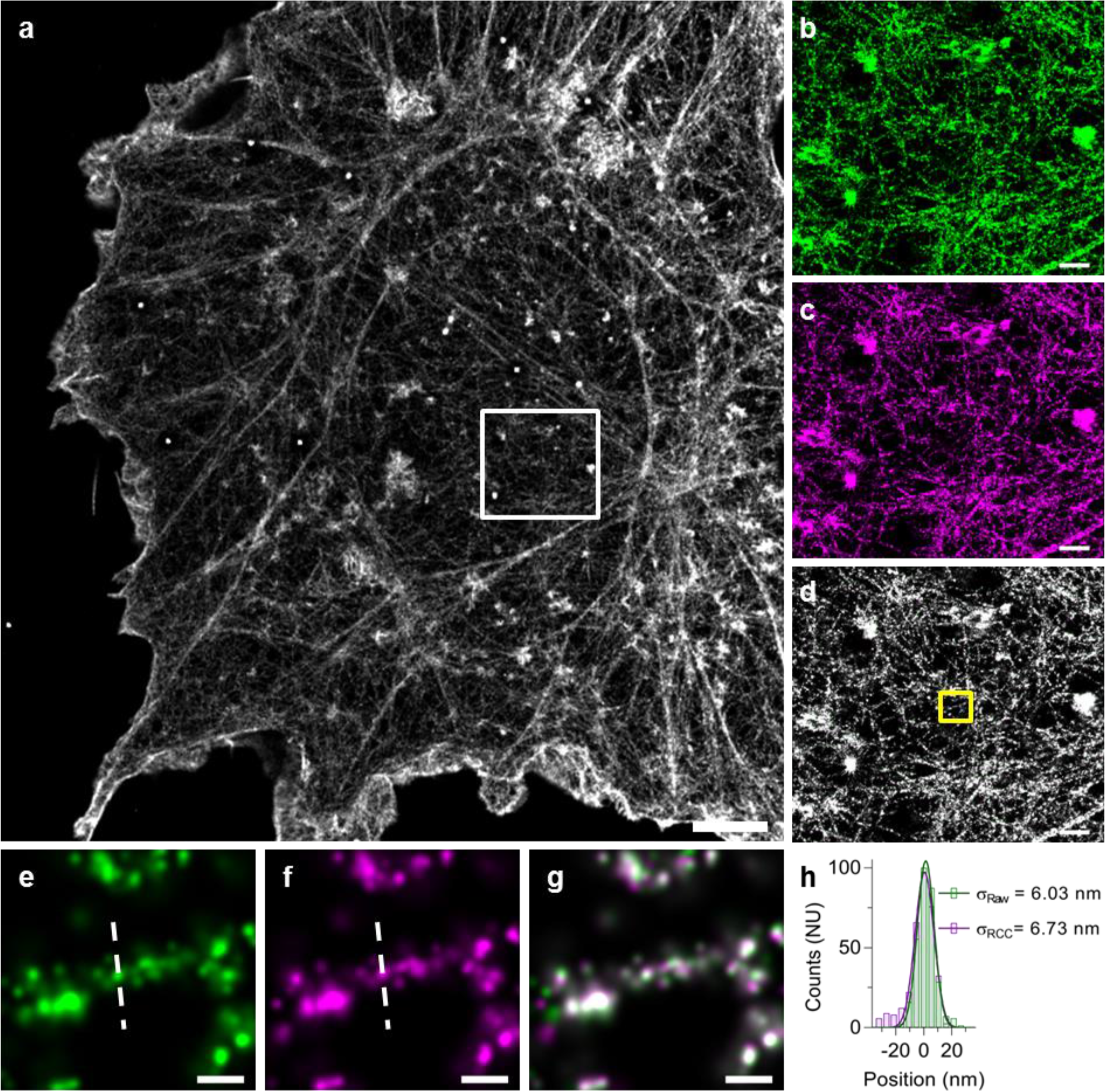
Post-acquisition drift correction does not improve the localization precision of Feedback SMLM data. (**a**) Raw data from the Feedback SMLM (green) and post-acquisition drift corrected data using RCC (magenta) were merged (white). Scale bar = 3 μm. (**b-d**) Zoomed region of the area (white square) highlighted in (a). Scale bars = 500 nm. (**e-g**) Image of selected, individual filaments (yellow square) indicated in (d). (**h**) Cross-sectional Gaussian profile fit of actin filament in (e) and (f). Scale bars = 50 nm.

**Supplementary Figure 13.**
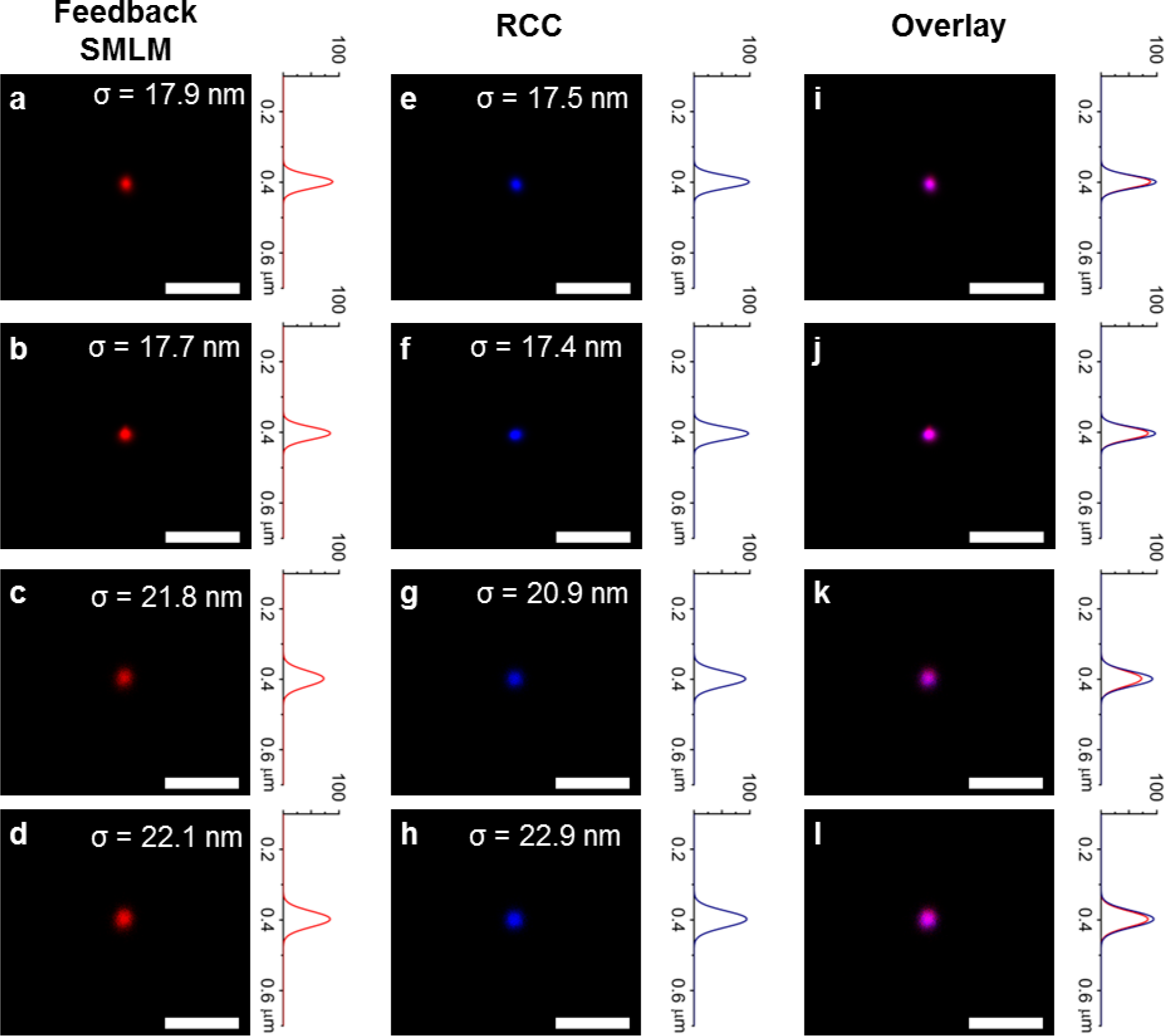
The Feedback SMLM does not exhibit residual drift. Gold nanorods were deposited onto glass coverslips to detect residual drift in Feedback SMLM. Performing post-acquisition drift correction did not enhance the resolution. (**a-d**) Raw data and distributions obtained with Feedback SMLM. (**e-h**) Data from (a-d) was drift corrected post-acquisition using RCC. (**i-l**) Data overlaid from (a-d) and (e-h). The distributions of the gold nanorods were fitted to Gaussian curves (a-l). Scale bar = 200 nm.

**Supplementary Figure 14.**
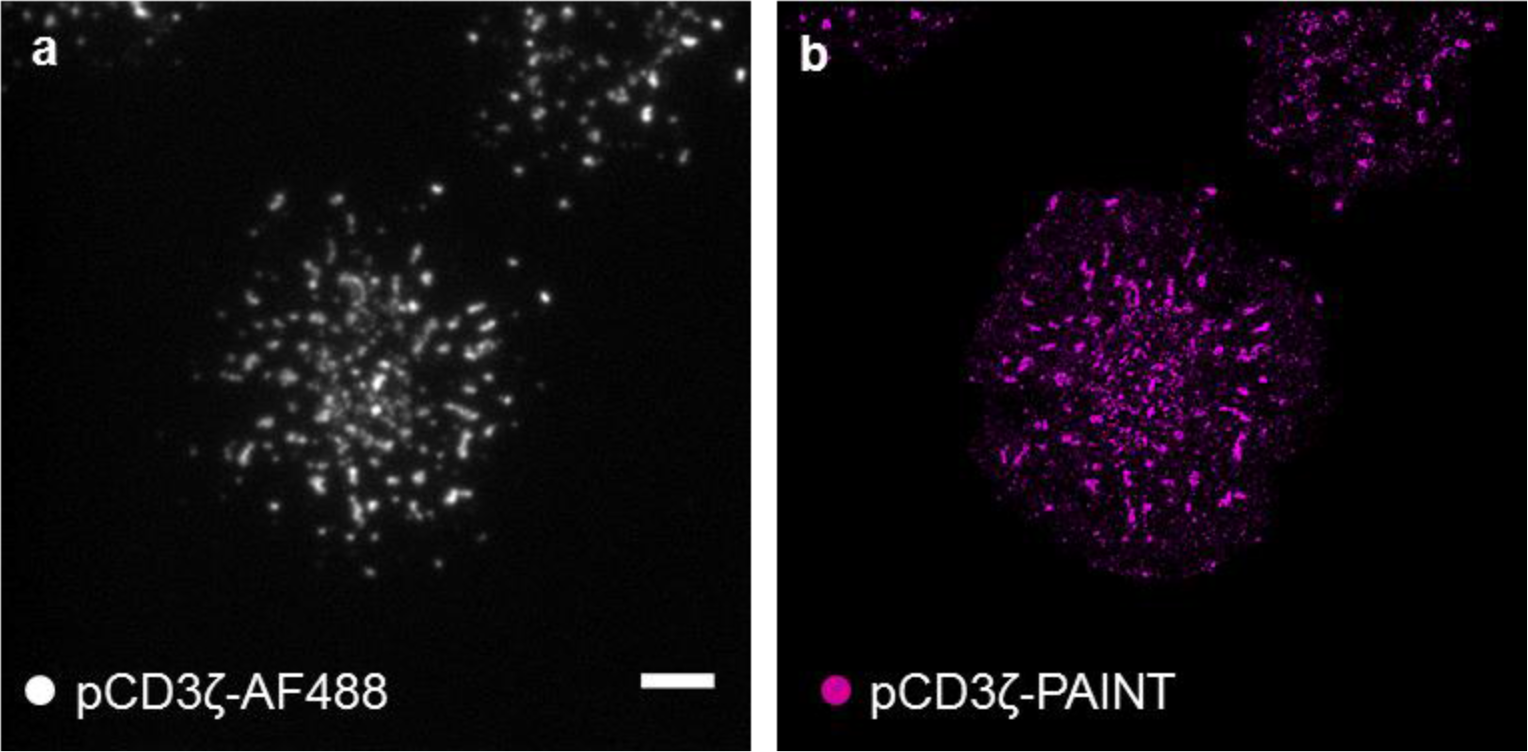
T cell activation. (**a**) TIRF image of antibody-stained phosphorylated CD3ζ. The secondary antibody was conjugated to Alexa 488. (**b**) DNA-PAINT image of phosphorylated CD3ζ (magenta) shows high spatial correlation with (A). Scale bar = 4 μm.

**Supplementary Figure 15.**
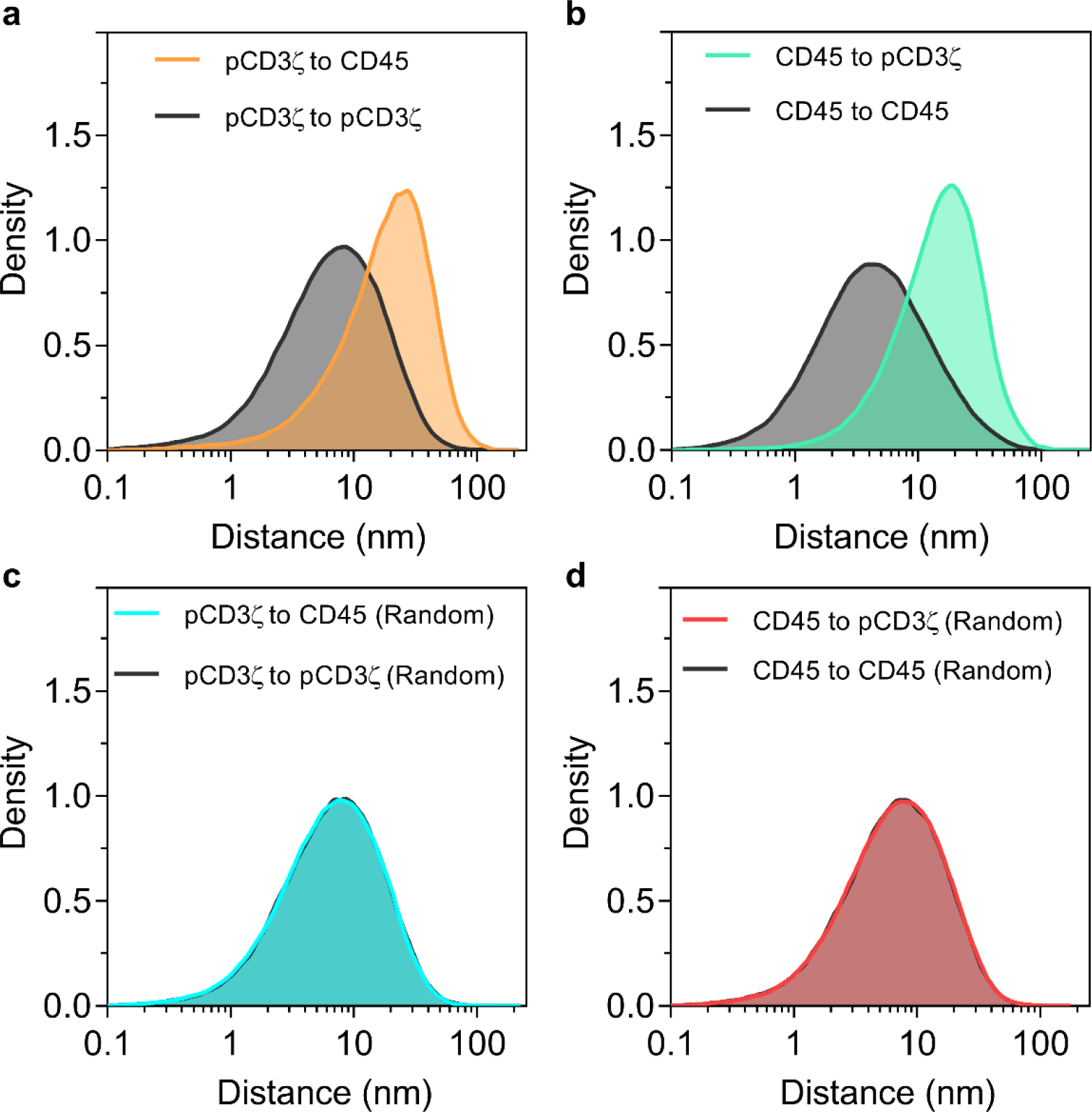
Nearest neighbor distance analysis over entire T cell. A T cell was activated on lipid bilayer with pMHC-I. Phosphorylated CD3ζ (pCD3ζ) and CD45 were imaged sequentially. Nearest neighbor distance (NND) analysis was performed on raw data between localizations of pCD3ζ and CD45. (**a**) NND of pCD3ζ to CD45 (orange line). NND of pCD3ζ to pCD3ζ localizations (black line) confirms spatial clustering of pCD3ζ and segregation from CD45. (**b**) NND of CD45 to pCD3ζ (green line). NND of CD45 to CD45 localizations (black line) confirms spatial clustering of CD45 and segregation from pCD3ζ. (**c** and **d**) A portion of randomly selected pCD3ζ and CD45 labels were exchanged and NND was performed. (**c**) NND of pCD3ζ to CD45 (blue line) and pCD3ζ to pCD3ζ (black line) using randomly exchanged labels. (**d**) NND of CD45 to pCD3ζ (red line) and CD45 to CD45 (black line) using randomly exchanged labels. The randomization underlines the specificity of the acquisition.

**Supplementary Figure 16.**
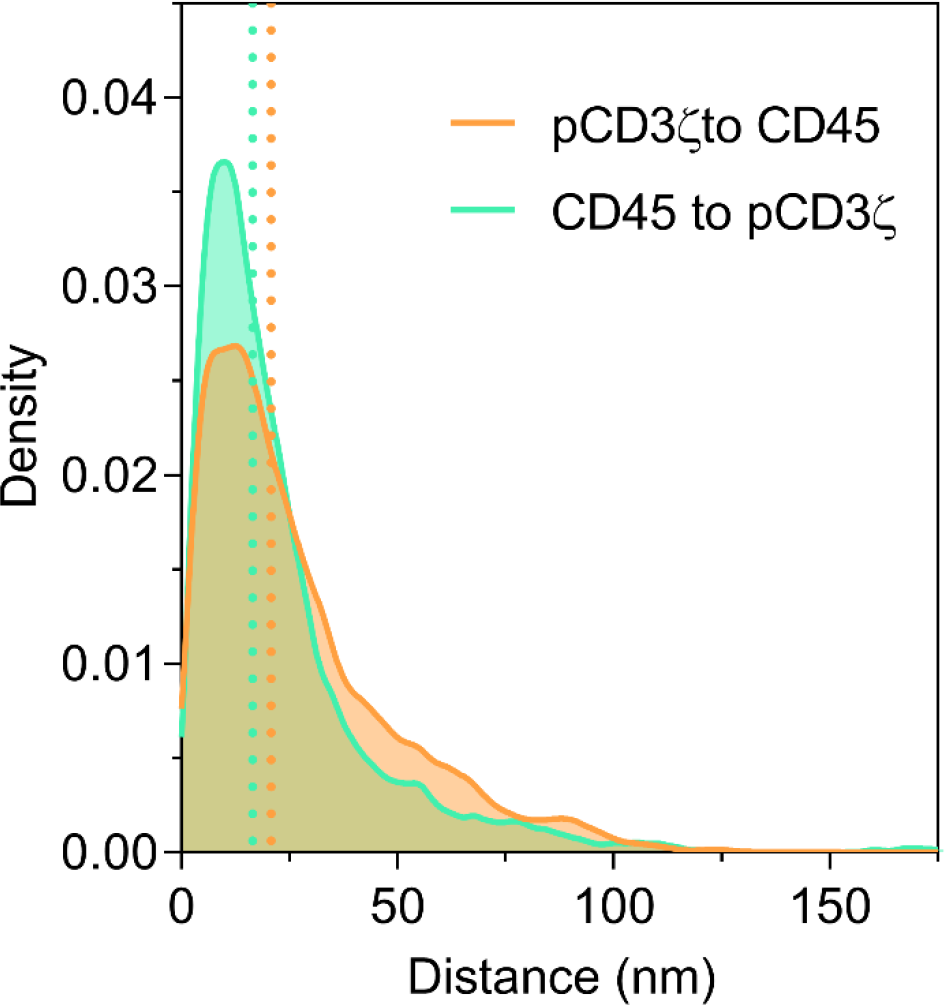
Nearest neighbor distance analysis of combined regions of interest between CD45 and pCD3ζ in Jurkat cells. Nearest neighbor distances between CD45 (green) and pCD3ζ (orange) for 40 individual regions of interest (10 per cell) were pooled. The median distance of pCD3ζ to CD45 is 20.8 nm (orange dotted line). The median distance of CD45 to pCD3ζ is 16.5 nm (green dotted line).

**Supplementary Figure 17.**
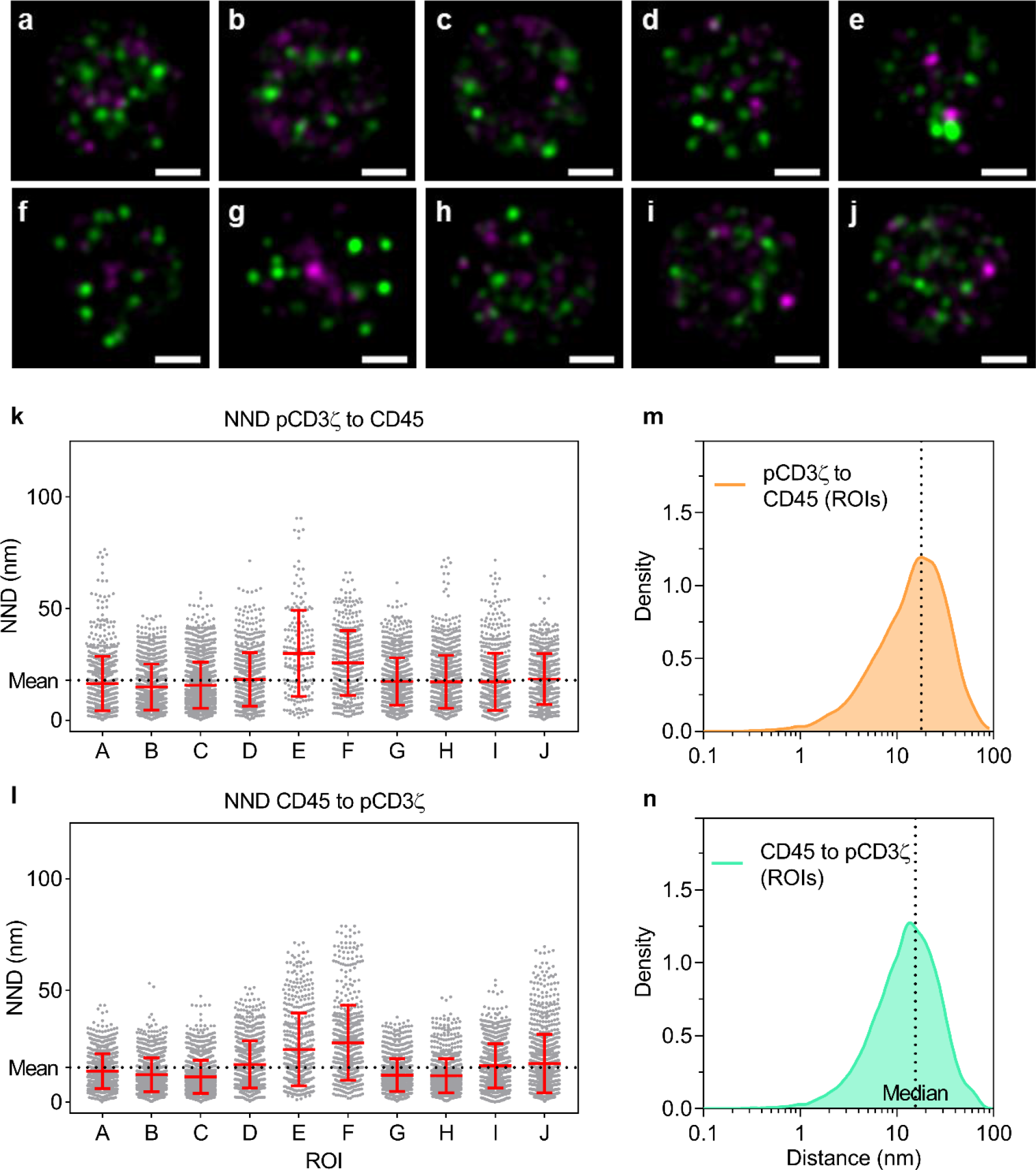
Nearest neighbor distance analysis of regions of interest between CD45 and pCD3ζ in Jurkat cells. (**a-j**) Individual regions of interest (ROI) used for nearest neighbor distance (NND), CD45 (green) and pCD3ζ (magenta) in Jurkat cells. Scale bars = 200 nm. (**k**) NND of pCD3ζ to CD45. (**l**) NND of CD45 to pCD3ζ. (k and l) Each symbol represents one distance measurement; horizontal and vertical bars represent the mean and standard deviation for each ROI. The mean across all ROIs is indicated by a dotted line. (**m**) NND distribution of pCD3ζ to CD45 (orange line) for the ROIs (A-J) with a median of 17.2 nm. (**n**) NND distribution of CD45 to pCD3ζ for the ROIs (a-j) with a median of 13.6 nm. (m and n) Dotted line represents the median distribution.

**Supplementary Figure 18.**
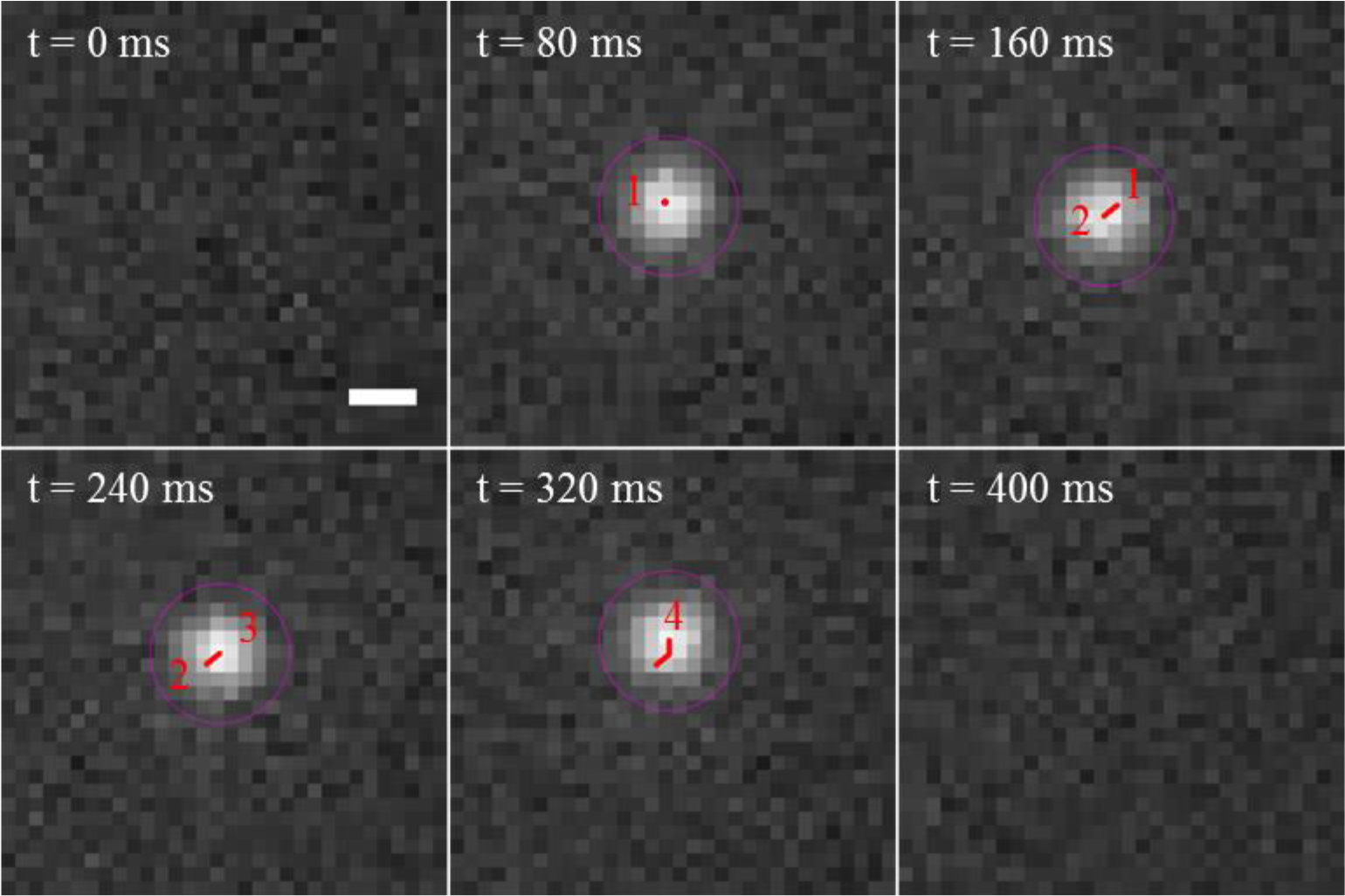
DNA origami structures are laterally mobile on lipid bilayers. DNA origami structures were deposited on lipid bilayers and the movement (red line, 1-4) of individual binding events (purple circle) monitored over time. Feedback SMLM could clearly detect the lateral movement. The movement was not due to drift as the sample was stabilized with a standard deviation of 0.39 nm and 1.1 nm in the lateral and axial directions, respectively. Scale bar = 400 nm.

**Supplementary Figure 19.**
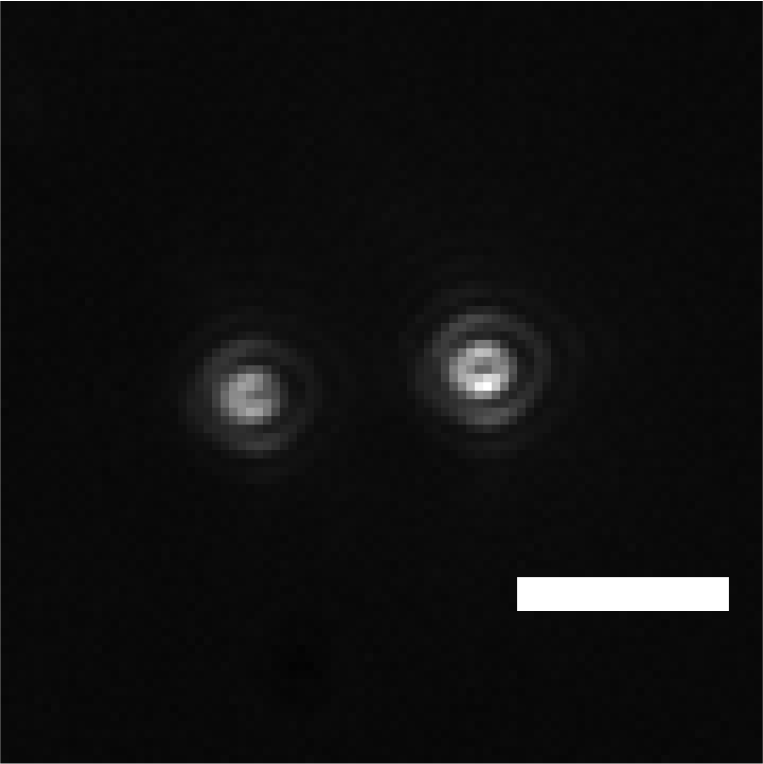
Gold nanorods can have orientation-dependent emission potentially impacting post-acquisition drift corrections. Gold nanorods were deposited on lipid bilayers and imaged. Once inserted into the bilayer, the emission profile detected is dependent on the orientation of the nanorod with respect to the laser. Typical post-acquisition drift correction software assumes fiducials are diffraction-limited Gaussian emitters and do not account for variations of the emission profile. The Feedback SMLM operates independently from the fluorescent acquisition and does not require post-acquisition correction. Scale bar = 2 μm.

**Supplementary Figure 20.**
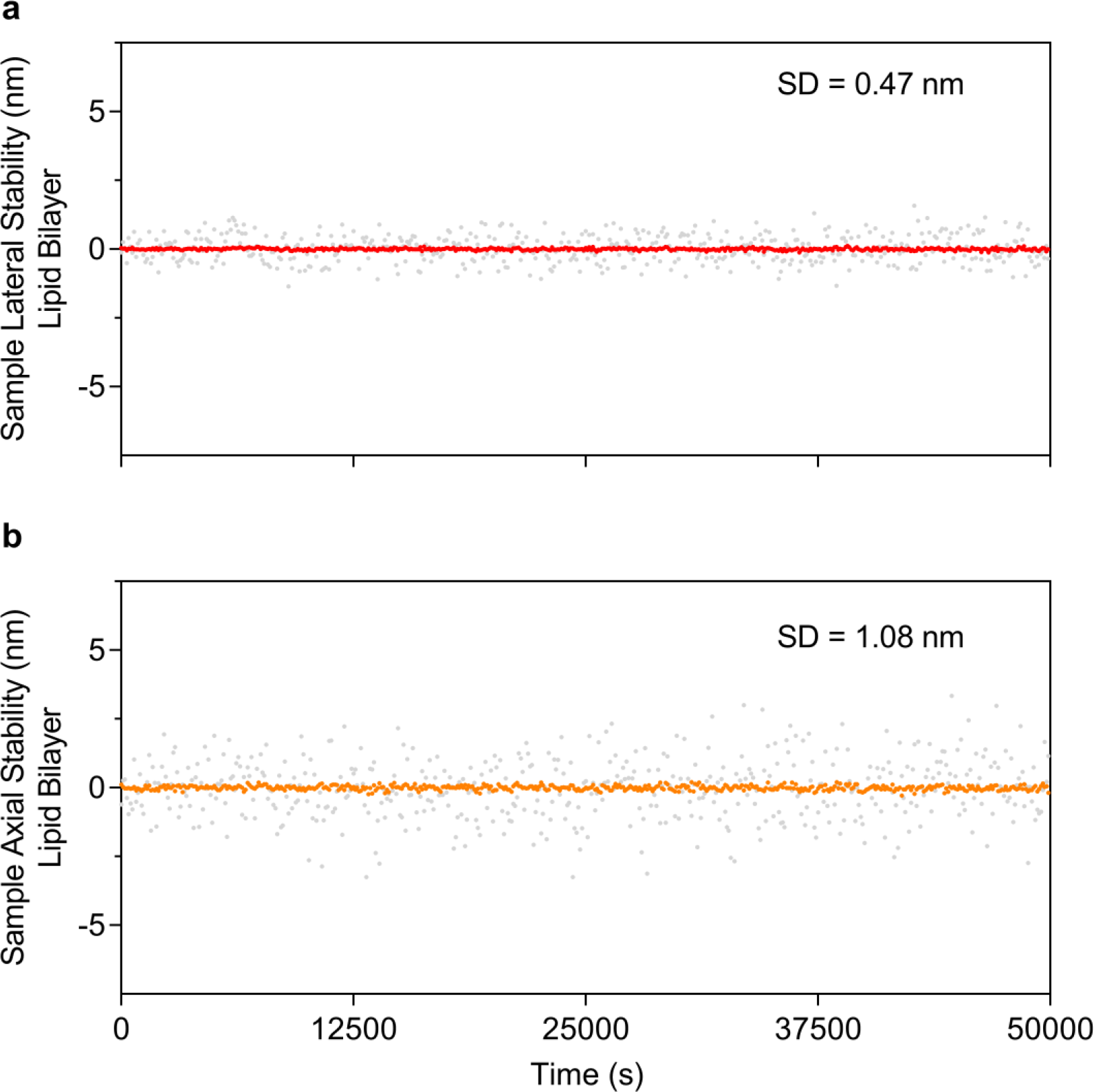
Sample stabilization on lipid bilayers. Beads were deposited on lipid bilayers with 1% biotin as described in the methods section. (**a**) Lateral stability of the sample with a standard deviation of 0.47 nm. (**b**) Axial stability of the sample with a standard deviation of 1.08 nm. Grey symbols represent sample deviation and red and orange lines represent a 10-point average (1/1000 points plotted).

**Supplementary Figure 21.**
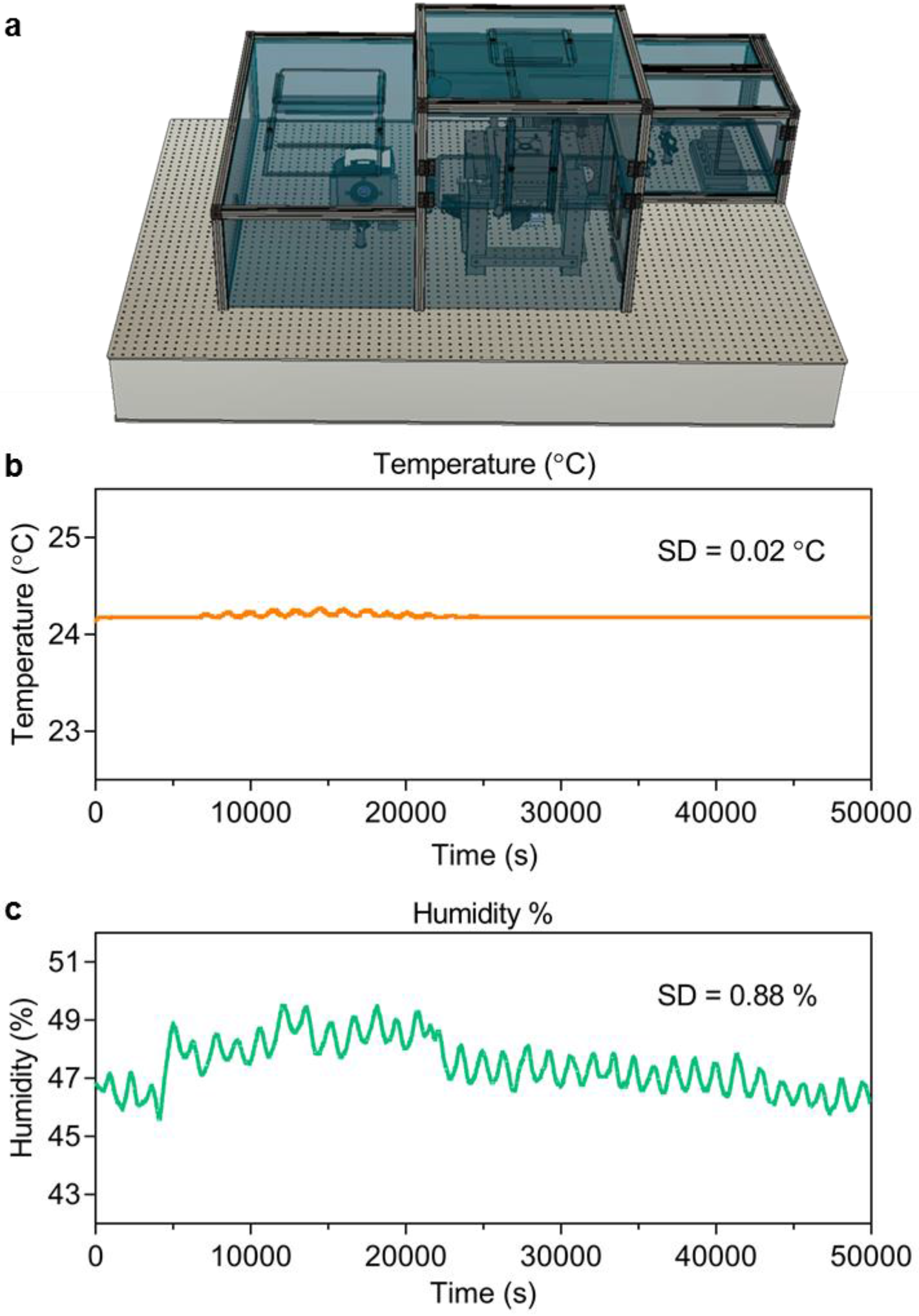
Environmental control. Subtle changes in temperature and humidity can induce vibrations and expansion of the optics within the microscope. (**a**) We developed an environmental chamber which compartmentalizes three different sections of the microscope and regulates each section individually. (**b**) The temperature was internally regulated with a standard deviation of 0.02°C. The temperature was typically set to +1°C above room temperature. A series of Peltiers were distributed throughout the system and regulated individually to counteract variations in room temperature. The temperature of the room has a standard deviation of ±0.5°C. (**c**) The humidity was regulated externally and monitored within the environmental chamber. Humidity has a standard deviation of 0.88%.

**Supplementary Table 1.** DNA-PAINT sequences.

| **Epitope** | **Docking Strand Sequence 5'-3'** | **Imaging Strand Sequence 5'-3'** | **Dye** | **Imaging strand name** | **Sequence reference** |
| --- | --- | --- | --- | --- | --- |
| **Gattaquant** | XX-Gattaquant | Gattaquant- XX - | ATTO 655 | G1 |  |
| **Phalloidin** | Biotin-<br>TTATACATCTA | CTAGATGTAT | ATTO 655 | IS_01 | Ref. 8 |
| **CD45** | TTTCTTCATTA | GTAATGAAGA | ATTO655N | IS_03 |  |
| **pCD3ζ** | TTATGTGAAAT | CATTTCACAT | ATTO655N | IS_22 |  |

**Supplementary Table 2.**
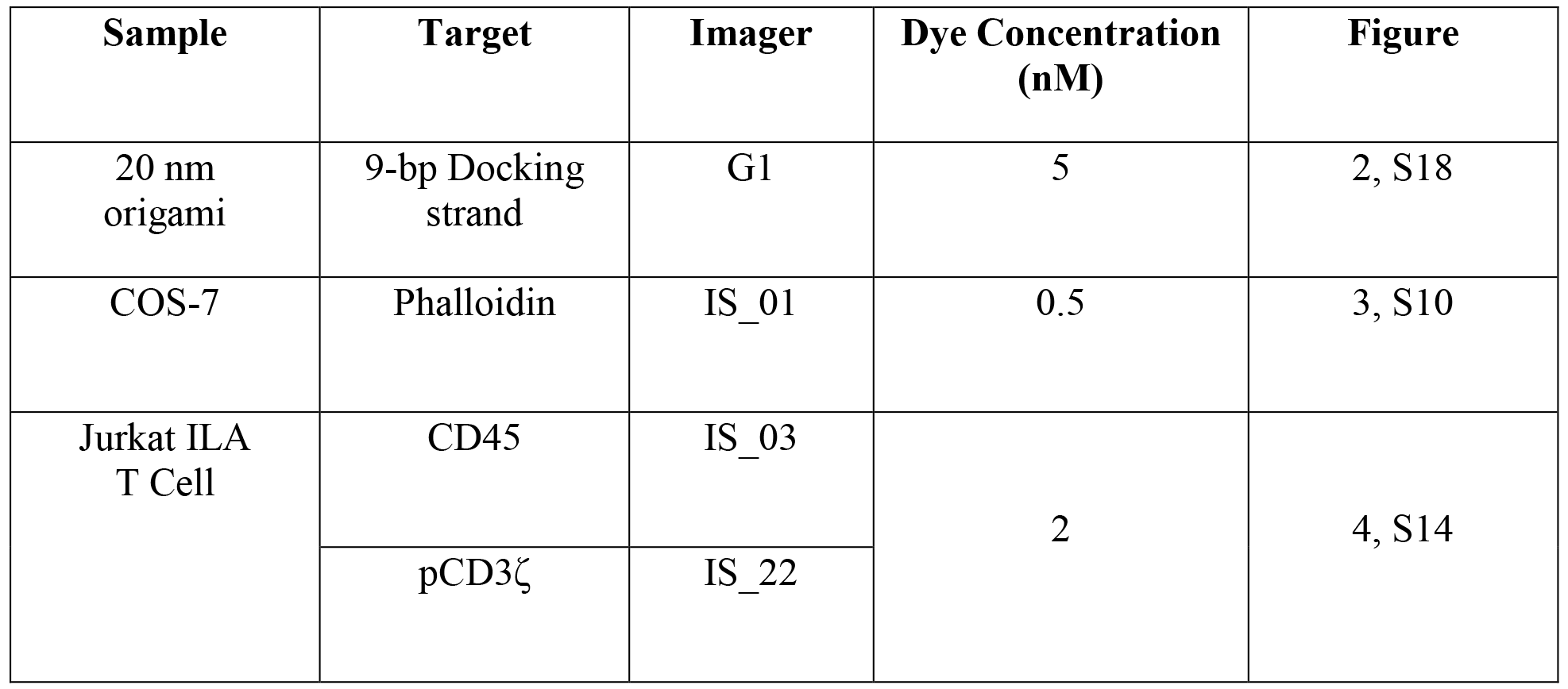
Sample parameters.

**Supplementary Table 3.** Fluorescence acquisition parameters.

| **Figure** | **Exposure time**<br>**(ms)** | **N Frames**<br>**(10<sup>3</sup>)** | **Gain** | **Laser Power**<br>**(kW/cm<sup>2</sup>)** |
| --- | --- | --- | --- | --- |
| 2 | 80 | 250 | 5 | 3 |
| 3 | 80 | 200 |  |  |
| 4, S14 | 200 | 200 |  |  |
| S7, S18, S19 | 80 | 1 |  |  |
| S10 | 80 | 150 |  |  |

**Supplementary Table 4.**
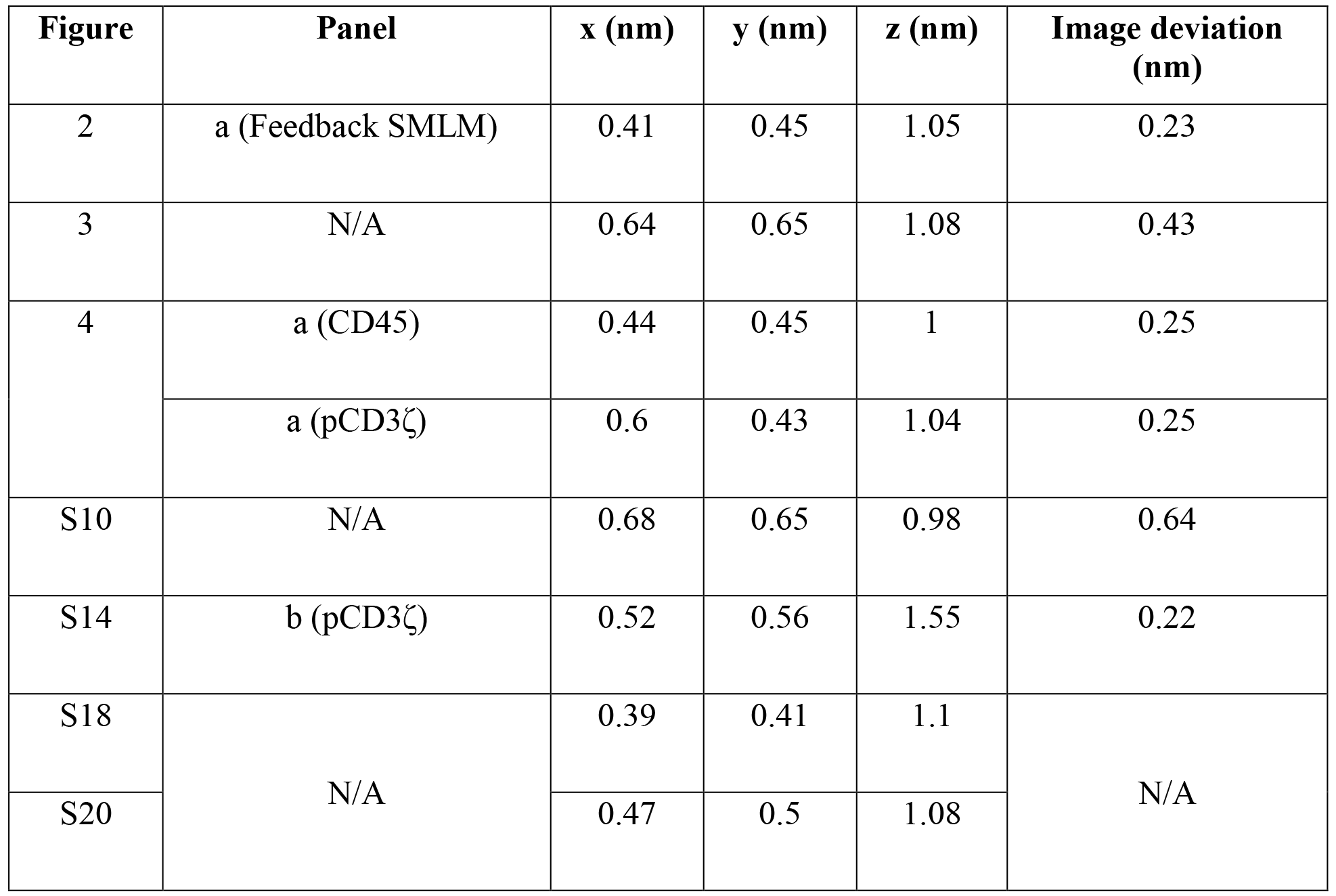
Sample stability.

| **Figure** | **Panel** | **x (nm)** | **y (nm)** | **z (nm)** | **Image deviation (nm)** |
| --- | --- | --- | --- | --- | --- |
| 2 | a (Feedback SMLM) | 0.41 | 0.45 | 1.05 | 0.23 |
| 3 | N/A | 0.64 | 0.65 | 1.08 | 0.43 |
| 4 | a (CD45) | 0.44 | 0.45 | 1 | 0.25 |
| | a (pCD3ζ) | 0.6 | 0.43 | 1.04 | 0.25 |
| S10 | N/A | 0.68 | 0.65 | 0.98 | 0.64 |
| S14 | b (pCD3ζ) | 0.52 | 0.56 | 1.55 | 0.22 |
| S18 | N/A | 0.39 | 0.41 | 1.1 | N/A |
| S20 |  | 0.47 | 0.5 | 1.08 |  |

